# Selective clonal persistence of human retroviruses in vivo: radial chromatin organization, integration site and host transcription

**DOI:** 10.1101/2021.11.10.467892

**Authors:** Anat Melamed, Tomas W Fitzgerald, Yuchuan Wang, Jian Ma, Ewan Birney, Charles R M Bangham

**Affiliations:** Department of Infectious Diseases, Faculty of Medicine, Imperial College London, London, United Kingdom; The European Bioinformatics Institute (EMBL-EBI), Cambridge, United Kingdom; Computational Biology Department, School of Computer Science, Carnegie Mellon University, Pittsburgh 15213, PA, USA

## Abstract

The human retroviruses HTLV-1 and HIV-1 persist in vivo, despite the host immune response and antiretroviral therapy, as a reservoir of latently infected T-cell clones. It is poorly understood what determines which clones survive in the reservoir and which are lost. We compared >160,000 HTLV-1 integration sites from T-cells isolated ex vivo from naturally-infected subjects with >230,000 integration sites from in vitro infection, to identify the genomic features that determine selective clonal survival. Three factors explained >40% of the observed variance in clone survival of HTLV-1 in vivo: the radial intranuclear position of the provirus, its absolute genomic distance from the centromere, and the intensity of host genome transcription flanking the provirus. The radial intranuclear position of the provirus and its distance from the centromere also explained ~7% of clonal persistence of HIV-1 in vivo. Selection for transcriptionally repressive nuclear compartments favours clonal persistence of human retroviruses in vivo.

## Introduction

Human T-cell leukemia virus type 1 (HTLV-1) persists in the host chiefly by clonal proliferation (*1*, *2*). A typical HTLV-1-infected host has 10^3^-10^6^ HTLV-1-infected T cell clones (*3*); each clone can be distinguished by the unique integration site of the single-copy provirus in the host genome (*4*). Every clone has its own characteristics of proviral expression, host gene expression, chromatin structure and equilibrium abundance; each of these attributes is influenced by the genomic integration site (*5*, *6*).

In primary infection, the initial virus spread is rapid (*7*). The proviral load (PVL, percentage of HTLV-1-infected peripheral blood mononuclear cells (PBMCs)) reaches an equilibrium or set-point in each host. The PVL can vary between hosts by over 1000-fold (*8*) and is proportional to the number of different HTLV-1^+^ T-cell clones (*9*). The PVL is partly determined by the host immune response (*10*); the force of selection exerted by the HTLV-1-specific cytotoxic T lymphocytes (CTLs) depends on the level of expression of HTLV-1 antigens (*11*). Both HTLV-1 and HIV-1 persist in a reservoir in vivo that depends partly on continued clonal proliferation (*1*, *12*, *13*)

We previously reported that, whereas initial integration of HTLV-1 shows no preference for any given chromosome, the HTLV-1^+^ clones that persist in vivo are found more often than by chance in the acrocentric chromosomes (13, 14, 15, 21 and 22) (*14*). The centromere-proximal regions of these chromosomes lie in the transcriptionally repressive environment around the nucleolus. This observation suggested (*14*) that repression of proviral expression in the nucleolar periphery minimizes the exposure of the infected cell to the strong anti-HTLV-1 immune response and so favours survival of that clone.

Chromosomes are not randomly distributed in the nucleus: each chromosome occupies a characteristic position known as a chromosome territory (CT) (*15*). The CTs are not static, but rather represent an average in the cell population. Both the CTs and individual chromosomes are radially organized in the nucleus (*16*): for example, chromosomes 18 and X often lie near the nuclear periphery, whereas chromosomes 17 and 19 are usually found in the centre of the nucleus (*17*–*19*).

The spatial distribution of transcriptional activity of the genome is also non-random. Intranuclear bodies known as nuclear speckles are associated with transcription and pre-mRNA processing (*20*, *21*). Two types of genomic domain are associated with particularly low transcriptional activity: lamina-associated domains (LADs) (*22*), near the nuclear periphery, and nucleolus-associated domains (NADs) (*23*–*25*). These domains are characterised by a relatively low gene density, a high density of repressive histone marks (H3K9me2/H3K9me3 in particular), and low GC content (*26*). There is a strong overlap between LADs and NADs, and certain domains stochastically re-associate with either the nuclear lamina or nucleolus in the daughter cells after mitosis (*24*, *27*).

The selective persistence of HTLV-1 proviruses in certain chromosomes, and the persistence in vivo of very different numbers of HTLV-1^+^ T cell clones in different hosts, imply that HTLV-1 infection results in the selective survival of certain clones, which then persist for the remainder of the host’s life. The aim of this study was to test two hypotheses that arise from these observations. First, that specific genomic attributes of the proviral integration site in an HTLV-1-infected clone determine its survival in the host during chronic infection. Second, that the intranuclear position of the provirus determines clonal survival in vivo. Finally, we applied the same approach to analyse data on HIV-1 integration sites.

## Results

### Analysis of integration sites identifies genome-wide correlates of clone survival

To identify the features of the HTLV-1 proviral integration site associated with clonal persistence in vivo, we compared specific genomic attributes between the integration sites isolated from PBMCs with those resulting from in vitro infection, which have not been subject to selection in vivo, in particular immune-mediated selection against virus-expressing cells. We compiled the data on HTLV-1 proviral integration sites from previous studies in our group (Table S1) into 3 unique datasets (Table 1): (1) in vivo sites - sites identified in PBMCs from naturally-infected individuals; (2) in vitro sites - collected from cells infected in culture; and (3) random sites - generated in silico by random selection of positions from a reference genome. All data were re-extracted from raw sequencing data (or mock data in the case of the random in silico sites) using the same pipeline, to ensure consistency of processing and to avoid systematic bias due to variable mapping quality.

**Table 1:** Datasets used in analysis

| Virus | Dataset type | Integration sites |
| --- | --- | --- |
| HTLV-1 | In vitro | 234607 |
| HTLV-1 | In vivo | 162401 |
| HIV-1 | In vitro | 65924 |
| HIV-1 | In vivo | 44367 |
|  | Random | 108976 |

### Preferential survival of HTLV-1 and HIV-1 is non-randomly distributed between and within chromosomes

While initial integration (in vitro) of HTLV-1 occurs in proportion to chromosome size (Figure S1; (*14*)), analysis of the much larger datasets in the present study confirmed the previous observation (*14*) of preferential survival of HTLV-1 in vivo in acrocentric chromosomes as a group. On closer inspection, this preference is seen to be due to a strong bias for survival in chromosomes 13, 14, and 15; no survival bias was observed in chromosome 21, and survival was counterselected in chromosome 22, the smallest acrocentric chromosome. The chromosome most preferred for survival in vivo is chromosome 18, and the chromosome most disfavoured for survival in vivo is chromosome 19 (Fig. 1A, Fig. S1). By contrast, in HIV-1-infected cells there was a stronger bias for or against initial integration in particular chromosomes than in HTLV-1-infected cells: the gene-rich chromosome 19 was the most favoured for initial HIV-1 integration, whereas the similarly-sized, gene-poor chromosome 18 was disfavoured (Fig. S2A-B). However, the rank order of chromosomes preferred for integration in vitro was very similar between HTLV-1 and HIV-1 (Kendall’s tau = 0.53, P < 0.001; Fig. S2C). The rank order of chromosome preference for in vivo survival was also correlated between the two viruses, albeit less strongly (Kendall’s tau = 0.32, P < 0.05; Fig. S2D).

**Fig. 1.**
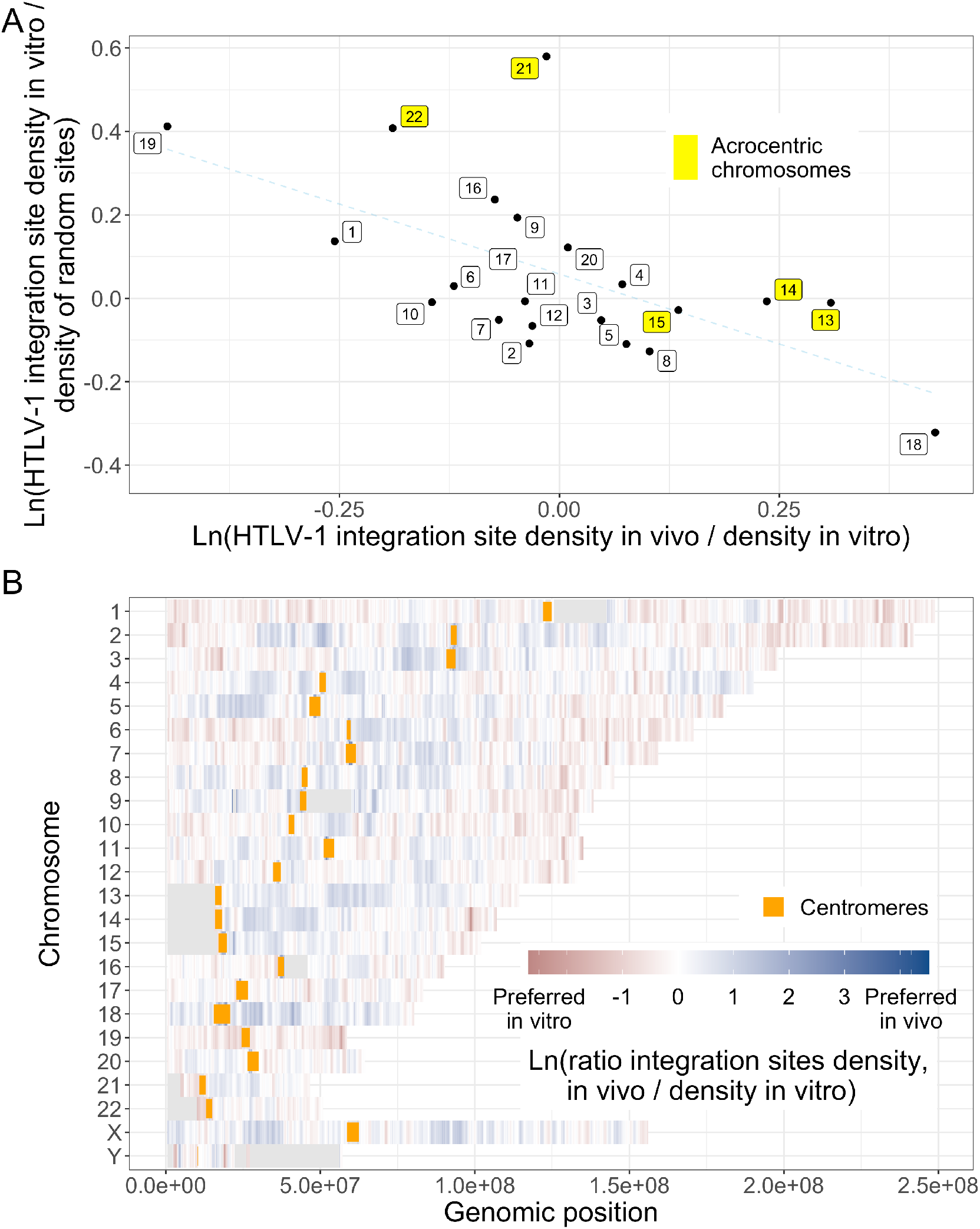
HTLV-1 Integration site survival is biased to specific chromosomes and is non-randomly distributed in each chromosome. (A) For each chromosome, two ratios of integration site frequencies (F) were calculated: F_in vivo_ / F_in vitro_ and F_in vitro_ / F_random sites_ (logarithmic scales). HTLV-1 survival is most strongly favoured in chromosome 18. The least-squares regression line is shown (dotted). (B) Heat-map of F_in vivo_ / F_in vitro_ in fixed windows across each chromosome (1 Mb wide, 1 kb steps): blue indicates preferential clone survival in vivo, and red indicates counterselection in vivo.

Within each chromosome, certain regions are either favoured or disfavoured for HTLV-1 survival in vivo. On most chromosomes, areas close to the centromere are more strongly favoured, whereas areas more distant from the centromere and closer to the telomere are disfavoured in vivo, in contrast with the more uniform distribution of initial integration in vitro (Fig. 1B).

### HTLV-1 initial integration favours accessible, active chromatin

Previous work has shown that initial integration of HTLV-1 was strongly preferred in close proximity to specific transcription factor binding sites (TFBS) (*5*). To extend this observation to additional TFBS, we used all transcription factor ChIP-Seq datasets published by the ENCODE project (*28*, *29*), using the B-cell line GM12878, comprising 156 ChIP-seq datasets, from 135 transcription factors (an equivalent dataset is not available on ENCODE for T-cells; Table S2). Each integration site (or random site as control) was annotated with respect to each TFBS dataset, and the minimum distance to any TFBS was calculated. The results show that integration sites were significantly enriched within 1 kb and 10 kb of any TFBS both in vivo and – more strongly – in vitro, compared with random sites. This observation suggests that initial integration is more frequent in accessible chromatin, available for transcription factor binding (Fig. S3A). To corroborate this conclusion, we identified the DNase I hypersensitive site (DHS) nearest to each integration site. We find that HTLV-1 sites within 10 kb of a DHS are more frequent than random expectation both in vivo and – especially – in vitro (Fig. 2A, Fig. S3B).

**Fig. 2.**
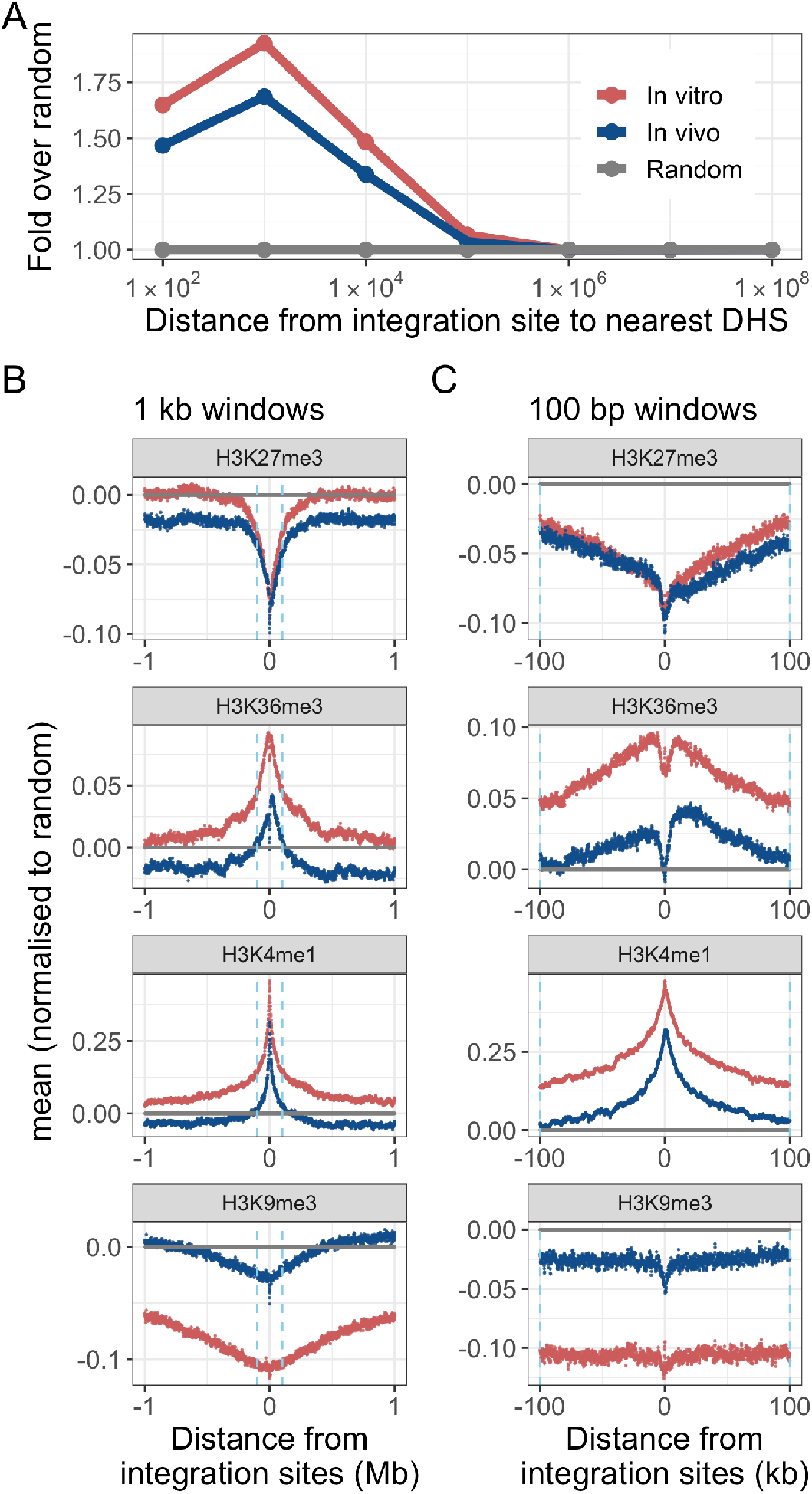
Initial HTLV-1 integration is biased towards accessible, transcriptionally active chromatin. HTLV-1 Integration site and random sites were mapped with respect to DNAse hypersensitive sites (DHS) and histone marks mapped by the ENCODE project using central memory T-cells. (A) Integration sites were enriched within 10 kb of a DNAse hypersensitive site (DHS). In each analysis, integration site frequency is compared to a random distribution. (B),(C) The mean density of each of 4 histone marks flanking the integration sites was calculated in discrete windows upstream and downstream across all integration sites from each dataset, either (B) - 1 kb windows, up to 1 Mb from the integration site, or (C) - 100bp windows, up to 100 kb from the integration site. Ln(fold change from random) is shown. Dashed line denotes 100 kb upstream or downstream.

Lastly, we used histone mark datasets from the ENCODE project to annotate the histone mark density in fixed windows, either 1 kb windows up to 1 Mb either side of the integration site (Fig. 2B, Fig. S3C), or 100 bp windows up to 100 kb either side of the integration site (Fig. 2C, Fig. S3D). We then calculated the mean density of each respective histone mark flanking the integration site. We found that for all histone marks associated with gene expression or activation (e.g. H3K36me3, H3K4me1, H3K27ac), the mean signal for in vitro integration sites was higher than random around the integration site, a preference that often extended (e.g. H3K4me1) to over 1 Mb either side of the integration site. By contrast, in vitro integration sites were less frequent near histone marks associated with transcriptional repression, either as a localized effect within 100 kb of the integration site (e.g. H3K27me3), or as a more distant effect, beyond 100 kb of the integration site (e.g. H3K9me3). Integration sites in vivo were less frequent than in vitro near all histone marks evaluated except H3K9me3, suggesting counterselection in vivo of proviruses that lie near those marks. Whereas the histone mark distribution around initial (in vitro) integration sites was symmetrical, the corresponding distribution around in vivo sites was in some cases asymmetrical. For example, H3K27me3 appeared to be counterselected in vivo more strongly downstream of the integration site than upstream (Fig. 2C, Fig. S3D).

### HTLV-1 survival and position along the chromosome

To identify the within-chromosome features that favour or disfavour survival of integrated HTLV-1 proviruses in vivo, we divided the human genome (GRCh38, see Methods) into discrete 1 Mb windows. We define the HTLV-1 survival index in each window as *Ln*(*F_in vivo_* / *F_in vitro_*), where *F* denotes the respective frequency of integration sites in that window.

We define the distance from each window to the centromere as the absolute distance between the midpoint of each window and the midpoint of the centromere. The results show a strong negative correlation between the HTLV-1 survival index and the distance from the centromere (Fig. 3) on both the p and the q arms of the chromosomes (Pearson’s R = −0.26 and −0.36, respectively): survival is favoured when the provirus is integrated closer to the centromere and progressively disfavoured towards the telomeres (Fig. S4). Survival of the integrated provirus correlated significantly less strongly with the distance from the telomere than with the distance to the centromere (Fig. S5).

**Fig. 3.**
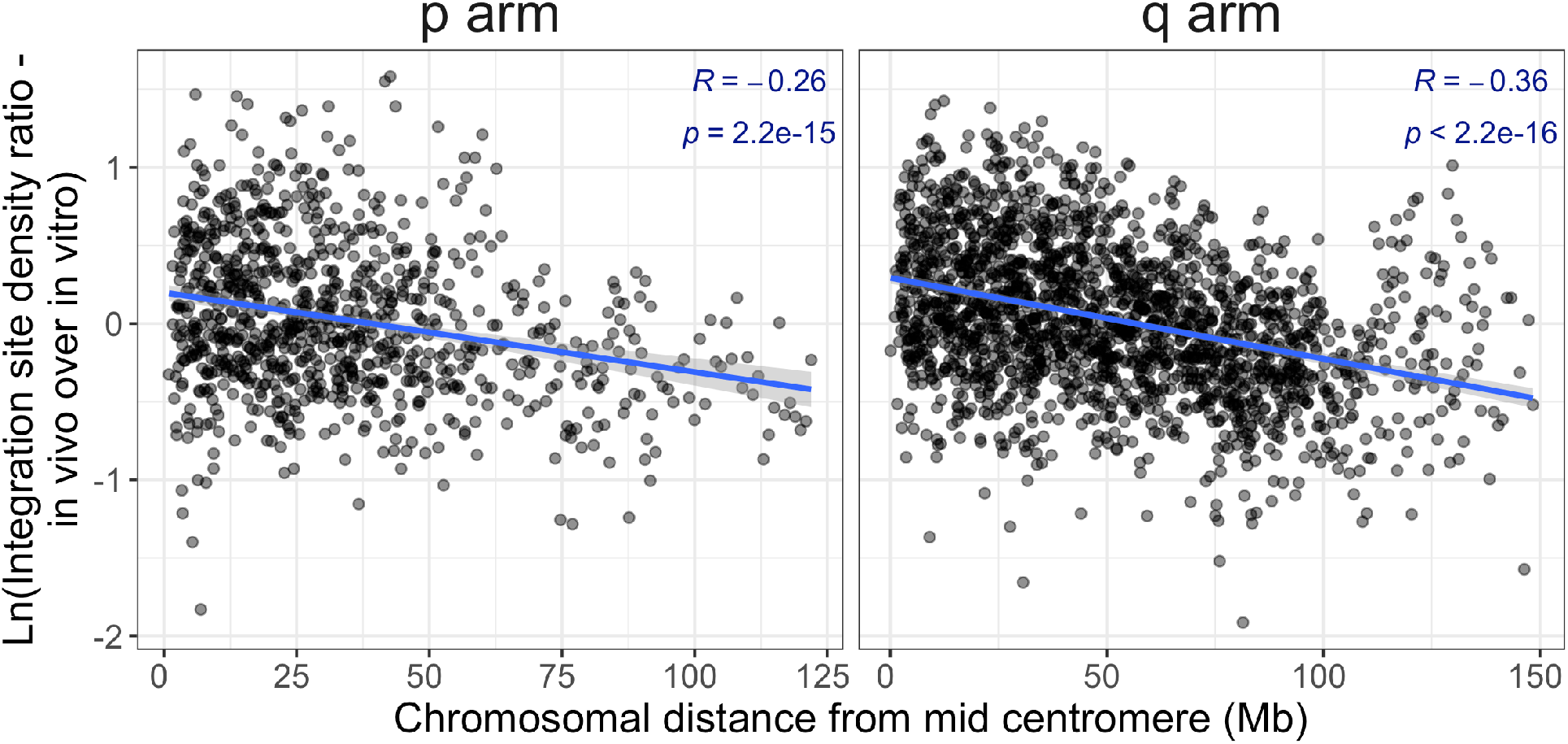
HTLV-1 survival vs distance from the centromere. The HTLV-1 clone survival index CSI_HTLV-1_ (defined as Ln(F_in vivo_ / F_in vitro_), the natural logarithm of the ratio between HTLV-1 in vivo and HTLV-1 in vitro site frequencies), is significantly negatively correlated with the absolute genomic distance from the centromere in both the short and long arms of the chromosomes (Pearson’s correlation test).

### HTLV-1 proviruses selectively survive in chromatin near the nuclear lamina and distant from nuclear speckles

The observed preferential survival of proviruses integrated in particular chromosomes (*14*, *30*) raised the question whether the physical position of the provirus in the nucleus influences HTLV-1 clonal survival in vivo. To answer this question, we estimated the distance of the provirus from specific intranuclear sites by using TSA-seq data published by Chen et al. (*20*) to estimate the distance of a given genomic location from the nuclear lamina or from nuclear speckles. We aligned and processed the integration site data according to the protocol described(*20*) and determined the mean TSA-seq signal in each 1 Mb window in which the integration site frequency was quantified. The results (Fig. 4A) showed a bias towards initial integration (in vitro) in chromatin that lies near nuclear speckles. By contrast, the HTLV-1 in vivo survival index showed a strong positive correlation with proximity to the lamin proteins, and a strong negative correlation with proximity to the SON protein (Fig. 4B, Fig. S6-S8). A similar analysis of HIV integration sites showed a marked preference towards integration near SON (Fig. S9), consistent with a recent report of frequent HIV-1 integration in nuclear speckle-associated domains (SPADs) (*31*). However, there was a trend towards increased survival away from nuclear speckles and near the lamin proteins, similar to that observed in HTLV-1 (Fig. 4C, Fig. S10).

**Fig. 4.**
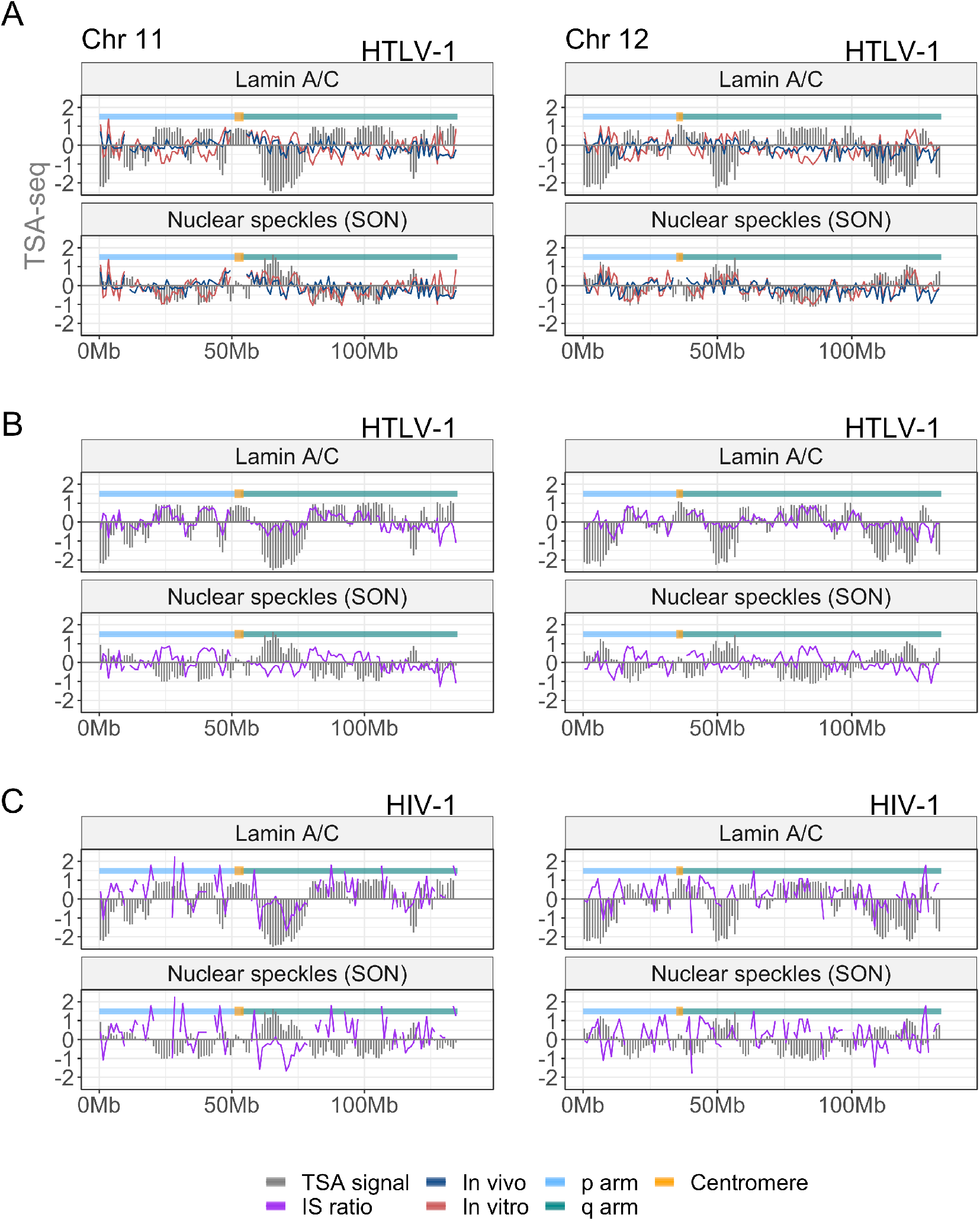
Preferential survival in vivo of infected T-cell clones whose provirus lies near the nuclear lamina and distant from nuclear speckles. Data on two mid-sized chromosomes (11 and 12) are shown; data on all chromosomes are shown in Supplementary Figures S6-S10. In each panel, the TSA-seq data on Lamin A/C and SON from (20) are plotted against Ln(integration site frequency, F). (A) HTLV-1 integration site frequency in vivo and in vitro. (B) The HTLV-1 clone survival index CSI_HTLV-1_ (Ln(F_in vivo_ / F_in vitro_)) closely tracks the Lamin A/C TSA-seq signal. (C) HIV-1 clone survival index, CSI_HIV-1_.

Genome-wide analysis confirmed a significant positive correlation between proximity to lamin proteins and the survival of HTLV-1 proviruses in vivo, and a significant negative correlation between proximity to nuclear speckles and survival (Fig. 5A, Fig. S11).

**Fig. 5.**
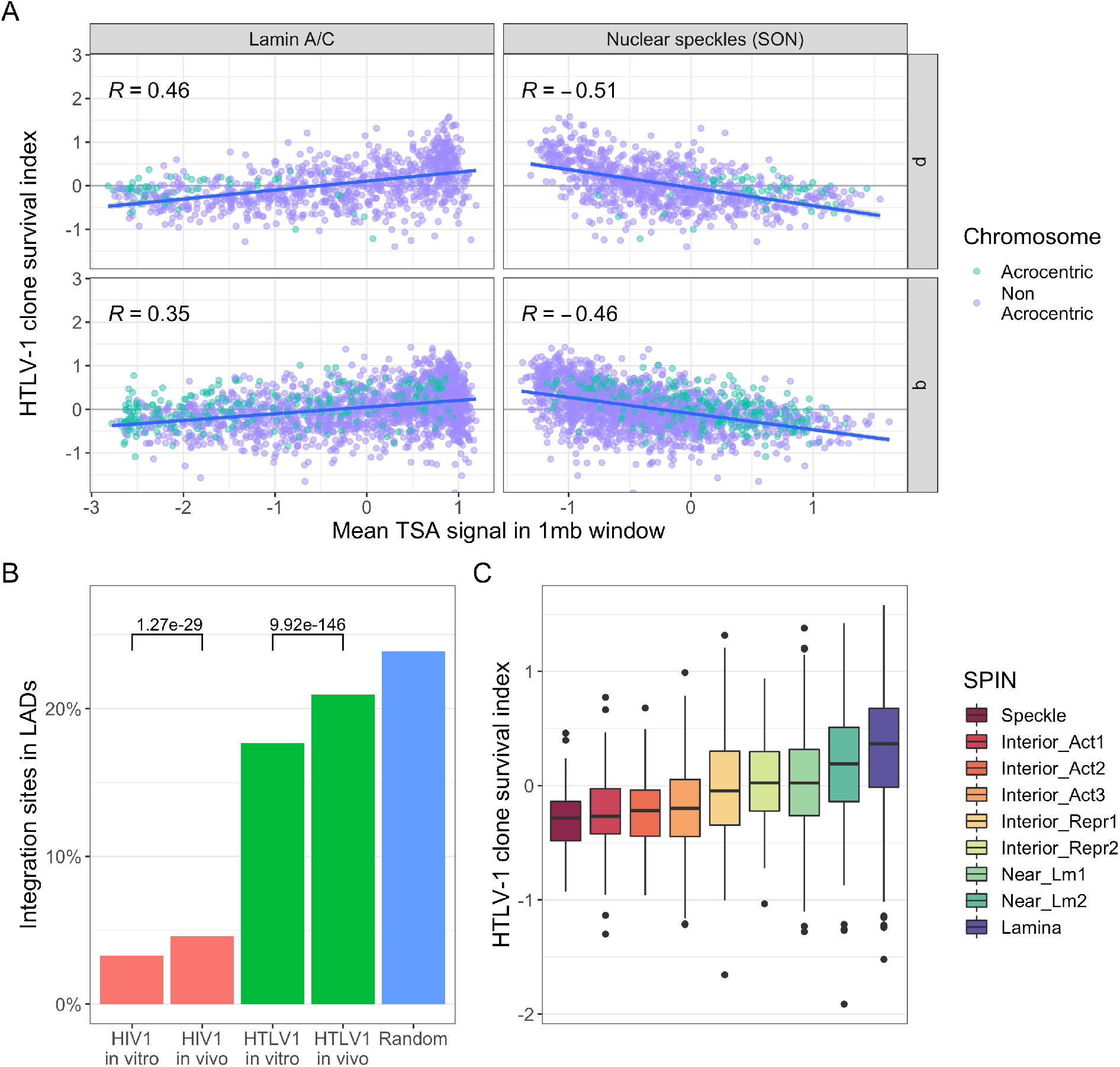
Genome-wide correlation between selective proviral survival and proximity to nuclear lamina, away from nuclear speckles. (A) TSA-seq data on lamin proteins and SON are significantly correlated with the clone survival index across the whole genome (Pearson’s correlation test, p < 10^−16^ for each test). Comparison with lamin B is shown in Figure S11. (B) Integration sites of both HTLV-1 and HIV-1 that lie near the nuclear lamina are significantly more frequent in vivo than in vitro. (C) SPIN state analysis (33) demonstrates a progressive increase in preferential survival of HTLV-1 towards the lamina.

Lamina-associated domains are usually identified using DamID. To corroborate the observation that proviral survival is associated with proximity to the nuclear lamina by an independent approach, we used a DamID dataset produced in T-cells (*32*). The results show that both in HTLV-1 and HIV-1, integration sites in vivo are enriched in lamina-associated domains compared to integration sites in vitro (Fig. 5B).

Lastly, we used the recently reported Spatial Position Inference of the Nuclear genome (SPIN) method, which combines data from TSA-seq, DamID and Hi-C to build a model which defines a set of spatial localization states of chromatin relative to nuclear bodies, reflecting a gradient of radial position from the nuclear lamina to the nuclear speckles (*33*). This analysis shows a monotonic increase in the HTLV-1 survival index towards the lamina (Fig. 5C).

### HTLV-1^+^ clone survival in vivo independently correlates with the expression status of the genomic region

The highest gene density is often located in “T” bands of the human genome, many of which are telomeric (*34*, *35*). We therefore investigated whether the observed decrease in survival associated with greater distance from the centromere could be attributed to an increase in gene density. In each 1 Mb window along the genome we quantified gene density using the Ensembl database and compared this density against the survival index. The results show that the gene density correlates with the distance to nuclear speckles (Fig. S12A-B), and is strongly negatively correlated with the survival index (Fig. S12C).

Because HTLV-1 is primarily found in vivo in CD4^+^ T-cells (*36*) we wished to test whether the expression of genes, specific to those cells, plays a role in the selective survival of integrated proviruses. Using expression data on primary T-cells from the Blueprint Epigenome project (*37*), we defined a mean expression level (regardless of the number or position of genes) in 1 Mb windows across the genome. In the observed bimodal distribution of expression intensity, we used the local minimum to define genomic windows which are low-expressing or high-expressing. We find that the HTLV-1 survival index is significantly lower in high-expressing genomic sites (p<10^−16^, Wilcoxon rank sum test). Similarly, at the level of 1 Mb windows, there was a strong negative correlation between expression intensity and the survival index (Fig. 6, Fig. S13).

**Fig. 6.**
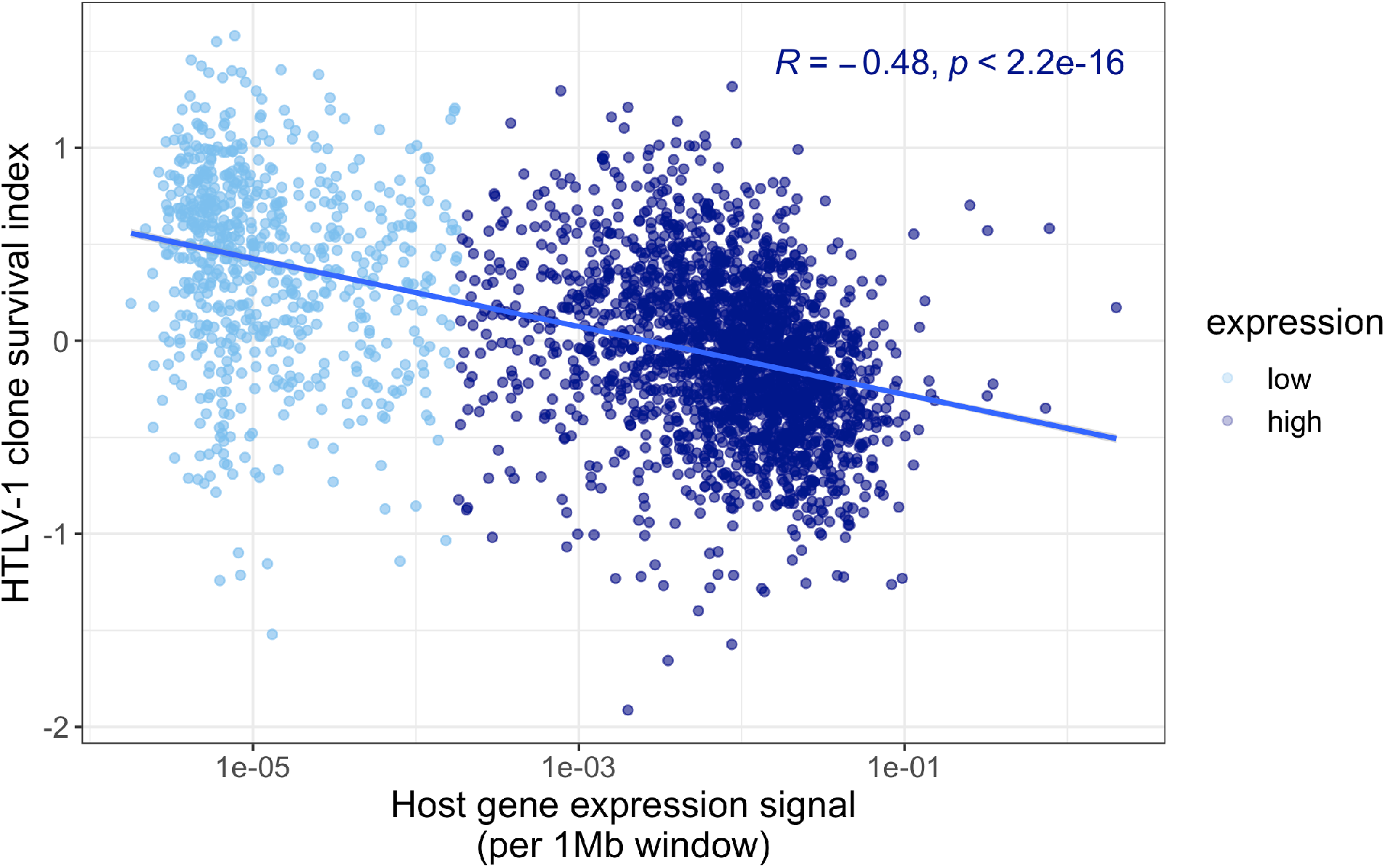
Counter-selection of HTLV-1 proviruses in highly expressing genomic regions. The HTLV-1 clone survival index CSI_HTLV-1_ (defined as Ln(F_in vivo_ / F_in vitro_), the natural logarithm of the ratio between HTLV-1 in vivo and HTLV-1 in vitro site frequencies), is significantly negatively correlated with the expression intensity in CD4+ T-cells (Pearson’s correlation test).

Since several genomic features considered here are known to be correlated, for example mean expression intensity and distance from the centromere, we carried out multivariate linear regression to identify the independent correlates of survival of HTLV-1^+^ clones in vivo. Three factors - distance to nuclear speckles (TSA-seq signal), the expression intensity and the distance to the centromere - remain significant independent predictors, together explaining ~40% of the observed variation in the HTLV-1 survival index (Table 2, Table S3, Fig. S14). A similar analysis of the HIV-1 data identified two of these factors - the distance to nuclear speckles and distance to the centromere - as independent correlates, together explaining ~7% of the observed variation in HIV-1+ clone survival in vivo (Table 2, Table S4).

**Table 2:** Linear model, significant predictors of clone survival index (CSI).

| predictor | HTLV-1 | HIV-1 |
| --- | --- | --- |
| (Intercept) | -0.12 | 0.25 |
| Proximity $P_s$ to nuclear speckle<sup>a</sup> | -0.38 | -0.33 |
| Distance $D_c$ from centromere (Gb) | -4.36 | -1.78 |
| $P_s * D_c$ | 2.87 | NS |
| Local host expression intensity<sup>b</sup> | -0.04 | NS |
| **N (1 Mb windows)** | **2677** | **2135** |
| **$R^2$ adjusted** | **0.41** | **0.07** |

<sup>a</sup> - TSA-seq signal for SON <sup>b</sup> - $\ln(\text{expression signal in T-cells})$

## Discussion

Like other persistent viruses, HTLV-1 establishes an equilibrium between viral replication and the host immune response. HTLV-1 does this by two chief mechanisms. First, by replicating mainly by clonal proliferation of infected cells rather than by de novo infection, thus minimizing the need for viral antigen expression and consequent immune-mediated killing. Second, by expressing the proviral plus-strand (which encodes the most immunogenic viral antigens) in rare, self-limiting bursts (*38*, *39*). The resulting reservoir of long-lived HTLV-1 clones is very large: the proviral load frequently exceeds 10% of PBMCs in non-malignant HTLV-1 infection.

The optimal strategy of survival for persistence of HTLV-1 in vivo is therefore to minimize proviral expression during most of the lifetime of the infected cell, but to retain the ability to re-express the provirus in intense bursts, either to infect a new host or to create a new clone in the same host (*1*).

The characteristics of proviral expression differ from clone to clone, and appear to be determined largely by the proviral integration site (*5*, *6*). We therefore hypothesized that local features of the chromatin flanking the provirus, such as epigenetic modifications associated with transcriptional activity, would correlate with the selective clonal survival of HTLV-1^+^ cells in vivo.

The results presented here show that certain epigenetic marks are indeed associated with in vivo survival of an HTLV-1^+^ T cell clone; however, these effects are relatively weak. By contrast, we found a remarkably strong correlation between selective in vivo clone survival and three factors: the spatial position of the provirus in the nucleus; the transcriptional activity of the host genome flanking the provirus; and the absolute genomic distance between the provirus and the centromere. Together, these factors explain ~40% of the observed variation in HTLV-1^+^ clone survival. A clone whose provirus lies in a genomic region with a tendency to locate near the nuclear lamina is more likely to persist in vivo than one whose provirus occupies a central position in the nucleus.

We conclude that integration of an HTLV-1 provirus into a genomic region that typically occupies a transcriptionally repressive compartment in the nucleus - near the nuclear lamina or the nucleolar periphery - favours the survival of that clone in vivo. Initial proviral integration favours transcriptionally active, accessible regions of the genome (Fig. 4A; (*9*) (*5*)), but the results reported here show that proviruses in regions of high transcriptional activity are counterselected during the subsequent chronic infection. The importance of the spatial intranuclear position of the provirus in the in vivo clone survival of human retroviruses is summarized in the model in Fig. 7.

**Fig. 7.**
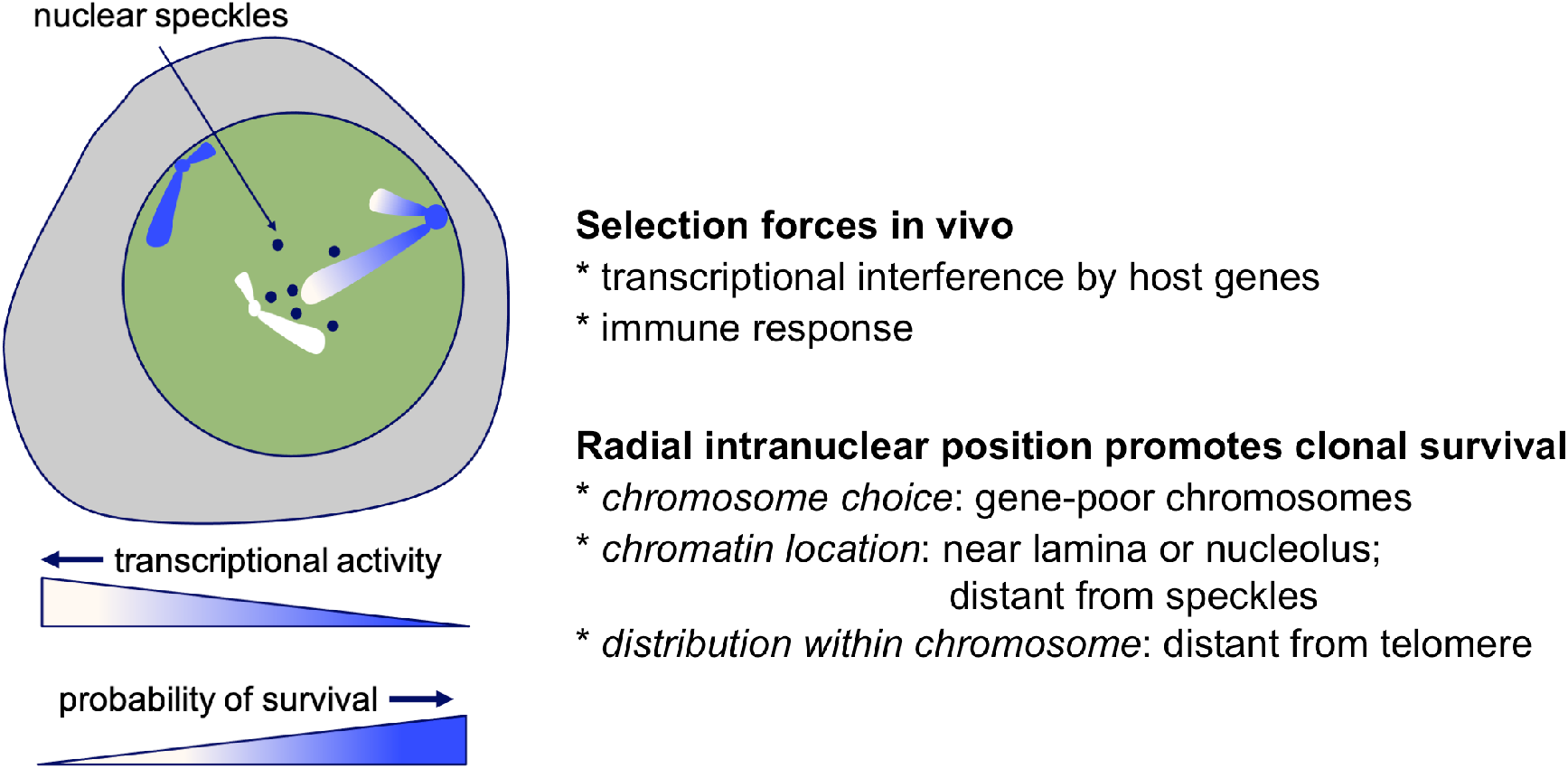
Proposed model of HTLV-1 selection in vivo. In this model, an infected T cell clone whose provirus is integrated in a genomic region that typically lies in a transcriptionally repressive compartment in the nucleus is more likely to survive in vivo in the face of immune-mediated selection, because the frequency of proviral reactivation is lower than that in regions of constitutively accessible chromatin, so the virus is less frequently exposed to the immune response. A clone whose provirus is integrated into constitutive heterochromatin is less likely to persist in vivo, because the clone lacks the proliferative advantage enjoyed by HTLV-1-expressing cells (1). Gene repression at the nuclear periphery and nucleolar periphery is not invariable (47); in addition, certain chromatin regions – facultative lamina-associated domains – are reversibly associated with the nuclear lamina (48). Thus, facultative heterochromatin may be the optimal site for in vivo survival, minimizing exposure to immune selection while retaining the potential to be re-expressed.

We previously showed that the HTLV-1 provirus binds the chromatin architectural protein CTCF (*40*), and thereby deregulates the higher-order structure and transcription of the flanking host genome (*6*). CTCF contributes to the localization of chromatin to the nucleolar periphery (*41*). It is therefore possible that CTCF binding provides an advantage to the virus by promoting association of the provirus with this transcriptionally repressive compartment.

HIV-1 differs strongly from HTLV-1 in its strategy of persistence in vivo. HTLV-1 expression is non-cytolytic, allowing clones to persist by intermittent proviral expression. By contrast, HIV-1 expression is cytolytic, and the virus persists in the host mainly by sustained de novo infection: that is, creation of new (albeit mostly short-lived) clones. However, the reservoir of HIV-1-infected cells that can persist indefinitely during highly active antiretroviral therapy is maintained partly by clonal proliferation (*12*, *13*) perhaps driven by normal homeostatic mechanisms (*42*). We applied the methods described above to analyse data on the HIV-1 proviral integration site, again from both in vitro infection and from cells isolated from infected individuals, both pre-ART and on-ART (*43*).

The results show that HIV-1 clone survival in vivo, like that of HTLV-1, is correlated with the nuclear position of the provirus (distance from nuclear speckles) and the distance from the centromere. This observation contrasts with the fact that the nuclear speckle-associated domains, which are enriched in transcriptionally active genes, are strongly favoured for the initial integration of HIV-1. However, in contrast with HTLV-1, only approximately 7% of the variation in proviral survival of HIV-1 can be explained by these factors: we postulate that this difference is due to the difference between the two retroviruses in the relative importance of infectious spread and mitotic spread (*1*) in the persistence of the virus in the host. The power of this analysis of HIV-1 data is limited by two factors. First, the smaller sample size (total of 110,338 integration sites of HIV-1, cf. 397,910 integration sites of HTLV-1). Second, the uncertainty in the proportion of HIV-1 integration sites identified in vivo that represent the true persistent reservoir, rather than short-lived clones. It is possible that the importance of the intranuclear spatial location of the HIV-1 provirus in persistence in vivo exceeds the estimate of 7% obtained here.

(*44*) examined the integration sites of HIV-1 in a group of elite controllers and individuals on antiretroviral therapy (number of genomes analysed = 1385 and 2388 respectively). They observed an overrepresentation of integration sites of genome-intact HIV-1 proviruses in centromeric satellite DNA, especially in elite controllers. These authors concluded that persistence of intact HIV-1 proviruses favours genomic integration sites that are transcriptionally silent or infrequently transcribed, and they suggested that the HIV-1 reservoir might resemble that of HTLV-1. Our conclusions are consistent with these observations and conclusions, and extend them by demonstrating in each virus the genome-wide importance of the position of the provirus both in the nucleus and within the chromosome.

In this study we used the data on the intranuclear distribution of chromatin from the TSA-seq analysis of K562 cells by (*20*) as well as the K562 SPIN states (*33*). K562 cells are an erythroleukemia line, and details of the intranuclear chromatin distribution in K562 may differ from that in T cells, which are the chief host cell infected by human retroviruses. Approximately 10% of the genome differs significantly between human cell lines in intranuclear position (relative to speckles) (*45*). The clonal survival of these human retroviruses may therefore correlate even more strongly with the spatial distribution of chromatin in T cells.

In HTLV-1 infection it is likely that the host immune response (*10*) is a major force that results in the observed pattern of selective clonal survival, by eliminating the clones that most frequently express the HTLV-1 provirus because the provirus is integrated either in a region of the host genome of intense transcriptional activity, or in a region that typically lies near the centre of the nucleus, near nuclear speckles and distant from the nuclear lamina and the nucleolus.

It is less clear why the absolute genomic distance between the provirus and the centromere is strongly correlated with survival, independently of the distance from nuclear speckles or the nuclear lamina. Average transcriptional intensity, gene density, GC content and early DNA replication all tend to increase towards the telomere. However, neither gene density nor GC content remains as a significant independent correlate of HTLV-1 survival in the multivariate regression analysis; and even after taking the transcriptional intensity into account, the distance from the centromere remains as a strong correlate of survival. The functional importance of DNA replication timing is not well understood (*46*). Our results suggest that HTLV-1 exploits some other, unidentified feature, independent of intranuclear position and transcriptional activity, that varies with the absolute genomic distance from the centromere.

## Methods

### Integration site datasets, cells and patients

The HTLV-1 integration site datasets used here (either in vivo or in vitro) are detailed in Table S1. Raw fastq data were used and processed in parallel to ensure consistency and comparability of data.

Fastq files were filtered to exclude potential spurious mapping events by selecting sequences that contain the final 5 bases of the HTLV-1 LTR (*49*)). Filtered sequencing reads were trimmed using trim galore (https://www.bioinformatics.babraham.ac.uk/projects/trim_galore/) to remove low quality and adapter bases, and subsequently aligned against a combined reference of hg38 human genome and HTLV-1 upstream sequence using BWA (*50*). Aligned reads were filtered using samtools (*51*) to include only uniquely mapped proper read pairs. Read pairs were further processed using a bespoke R script to correct the mapped position based on the CIGAR string and grouped based on unique pairs between integration sites and shear sites. Lastly, integration site abundance was estimated using the R package sonicLength (*52*) and cleaned to correct for mapping and barcode errors.

In vitro integration sites and donor cell line (MT-2) integration sites were sequenced in parallel. Any integration sites found in the donor cell line or in >1 infection assay were excluded from analysis.

Random sites were selected from the hg38 genome reference using the R package intSiteRetriever (*53*), and a mock fastq file was generated from these positions and hg38 sequence to simulate integration site raw data. Subsequently, this mock fastq file was processed through the same pipeline described above to ensure compatibility with integration site data.

The three main types of integration site were combined (HTLV-1 in vivo sites, HTLV-1 in vitro sites, Random sites) and repeatedly observed sites were removed from each dataset, to ensure non-redundancy. See Table 1 for summary of integration site counts.

### Integration site frequency and clone survival index

For analysis in 1 Mb windows across the human genome, discrete windows along each chromosome were defined from position 1 to the chromosome terminus. 1 Mb windows that overlap the end of the chromosome were excluded. To improve mapping confidence, 1 Mb windows with at least 1kb overlap with an ENCODE exclusion list region (*54*), or which include at least 1000 ambiguous bases (counted using bedtools nuc (*55*)) in the hg38 reference were also excluded.

The proviral integration site frequency (F) is calculated in each defined genomic region (e.g. 1 Mb window) as the proportion of all integration sites of a given dataset present in that region. We define a clone survival index (CSI) for each specified genomic region (e.g. 1 Mb window) as:

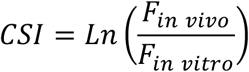

where CSI is undefined (either *F_in vivo_* or *F_in vitro_* = 0), the corresponding 1 Mb window is excluded from statistical analysis.

### HIV integration sites

We used data on HIV integration sites published by (*43*). In vitro (PHA-only) and in vivo (both pre-ART and post-ART) integration sites were compiled and remapped to hg38 using the liftover tool included in the R package rtracklayer (*56*). In keeping with the processing of the HTLV-1 data, if any two HIV-1 integration sites were mapped within 5 bp of each other (~1.2% of integration sites) one of the pair was removed to create a unique list of integration sites.

### Chromatin modification and accessibility annotation

To map the presence of integration sites with respect to histone mark density, transcription factor binding sites and DNAse hypersensitive sites, we used data from experiments carried out and analysed by the ENCODE project (*28*, *29*) (Table S2).

#### DNAse hypersensitive sites (DHS)

DNAse-seq data were retrieved from the ENCODE project using the following criteria: Organ - blood; Cell - leukocyte; Biosample - GM12878 or CD4-positive, alpha-beta memory T cell; genome assembly - GRCh38; genome assembly - GRCh38; filetype - “bed narrowpeak.” For GM12878 two replicate experiments are reported; a site is recorded as within N bases of a DHS if this condition is satisfied in both experimental replicates. Genomic distances are cumulative, e.g., the integration sites within 10 kb of a DHS also include the integration sites within 1 kb. Annotation of the nearest DHS to each integration site was done using the hiAnnotator R package (*57*).

#### Transcription factor binding sites (TFBS)

TF ChIP-seq datasets were retrieved from the ENCODE project using the following criteria: Organ - blood; Cell - leukocyte; Biosample - GM12878; genome assembly - GRCh38; filetype - “bed narrowpeak”; output type - “optimal idr threshold peaks”; Audit category excluding “extremely low read depth” and “extremely low read length.” At the time of retrieval (August 2019) 156 datasets were available from 135 targets. Where more than one dataset was available for the same target, the larger dataset was used. Annotation of the nearest TFBS to each integration site from each type was made using the hiAnnotator R package.

#### Histone mark data

Histone modification ChIP-seq datasets were retrieved from the ENCODE project using the following criteria: Organ - blood; Cell - leukocyte; Biosample - GM12878 or CD4-positive, alpha-beta memory T cell; genome assembly - GRCh38; filetype - “bigwig”; output type - “fold change over control.” Only those based on two replicates are used. Where more than one dataset was available for the same target, the larger dataset was used. Histone modification signal was averaged over fixed windows in the regions flanking each integration (or random) site using the UCSC bigWigAverageOverBed tool (*58*) and averaged across all integration sites.

### Nuclear position annotation

#### TSA-Seq data analysis

For consistency of reference (hg38), raw fastq data reported by (*20*) (Table S2) were realigned and processed according to the authors’ protocol (https://github.com/ma-compbio/TSA-Seq-toolkit). A Y-excluded reference genome (hg38F) was used (K562 is a female cell line); Bowtie2 (*59*) was used to align raw data + controls, followed by normalization using the authors’ script. Wig output was converted to BigWig format using UCSC BigToBigwig (*58*), which was then quantified in fixed 1 Mb windows across the human genome using UCSC bigWigAverageOverBed tool (*58*).

#### DAM-ID data preparation

Lamina-associated domain (LAD) data (Table S2) were mapped by (*32*) using the hg19 reference genome. LAD genomic positions were converted to hg38 using the liftover tool included in the R package rtracklayer (*56*). Integration sites in LADs were annotated using the hiAnnotator R package (*57*). Here, we show results from annotation against LADs in activated T-cells. Using the data from resting T-cells does not qualitatively alter these results.

#### Chromosome length

Chromosome positions, chromosome arms, gaps and centromere data were retrieved from the UCSC table browser (*60*).

### SPIN data analysis

We compared the HTLV-1 survival ratio on different SPIN states (*33*) identified in the K562 cell line. The SPIN states were calculated at 25 kb resolution and analyzed in the human hg38 genome assembly. First, we assigned SPIN states to each 1 Mb genomic bin. We used bedtools intersect (*55*) to calculate the overlap between SPIN states and predetermined 1 Mb genomic bins. We only kept 1 Mb genomic bins where the majority (≥ 75%) of regions were covered by a single SPIN state; genomic bins with less than 75% of regions registered with one single SPIN state were discarded. We then calculated the natural logarithm of the ratio of in vivo/in vitro proportion of integration sites (‘survival ratio’) for each 1 Mb bin. Genomic bins with zero integration or missing data were also discarded. Finally, we plotted the distribution of survival ratio on different SPIN states in the boxplot (Fig. 5C).

### Gene density and T-cell expression data

A gene position list for gene density quantification was retrieved from Ensembl BioMart database (version 80) using the biomaRt R package (*61*). The number of genes overlapping each 1 Mb window (overlaps of any length) was counted using the GenomicRanges R package (*62*).

For analysis of the local genomic expression signal, ribodepleted RNA-seq expression signal datasets were retrieved in bigwig format from the BLUEPRINT project site on http://dcc.blueprint-epigenome.eu/#/files based on the following criteria: Cell-type - central memory CD4-positive, alpha-beta T cell; tissue - venous blood. Using discrete 1 Mb windows across the human genome, the mean expression signal from uniquely mapped reads per window was calculated using the UCSC BigWigAverageOverBed tool (*58*), and the mean of available 2 samples calculated per window.

This study makes use of data generated by the BLUEPRINT Consortium. A full list of the investigators who contributed to the generation of the data is available from www.blueprint-epigenome.eu. Funding for the project was provided by the European Union’s Seventh Framework Programme (FP7/2007-2013) under grant agreement no 282510 - BLUEPRINT.

## Acknowledgments

The authors thank Andrew Belmont, Wendy Bickmore, Angus Lamond, Brian McStay, and members of the Bangham group, for helpful discussions. We thank Graham Taylor, Lucy Cook, the donors and research nurses in the National Centre for Human Retrovirology, Imperial College London; and Laurence Game and colleagues in the Genomics Facility in the MRC London Institute for Medical Sciences, London, UK. A large dataset of in vitro integration sites of HTLV-1 was obtained by Amy McCallin; we thank Goedele Maertens and Amy for these data.

## Author contributions

Conceptualization: AM, EB, CRMB; Data curation: AM; Software: AM, TWF, YW, JM; Formal analysis: AM, TWF, YW; Validation: AM, TWF; Investigation: AM, TWF; Visualization: AM, YW; Methodology: AM, TWF, EB, CRMB; Writing— original draft: AM, YW, CRMB; Project administration: AM, CRMB; Resources: CRMB; Supervision: JM, EB, CRMB; Funding acquisition: CRMB; Writing—review and editing: AM, TWF, JM, EB, CRMB.

## Data and materials availability

All data are available in the main text or the supplementary materials.

