## Supplemental Figures and Tables for "Selective clonal persistence of human retroviruses in vivo: radial chromatin organization, integration site and host transcription"

### Supplementary Materials

Anat Melamed*, Tomas W Fitzgerald, Yuchuan Wang, Jian Ma, Ewan Birney and Charles R M Bangham*

**This PDF file includes:**

Figs. S1 to S14

Tables S1 to S4

**Other Supplementary Materials for this manuscript include the following:**

Data S1

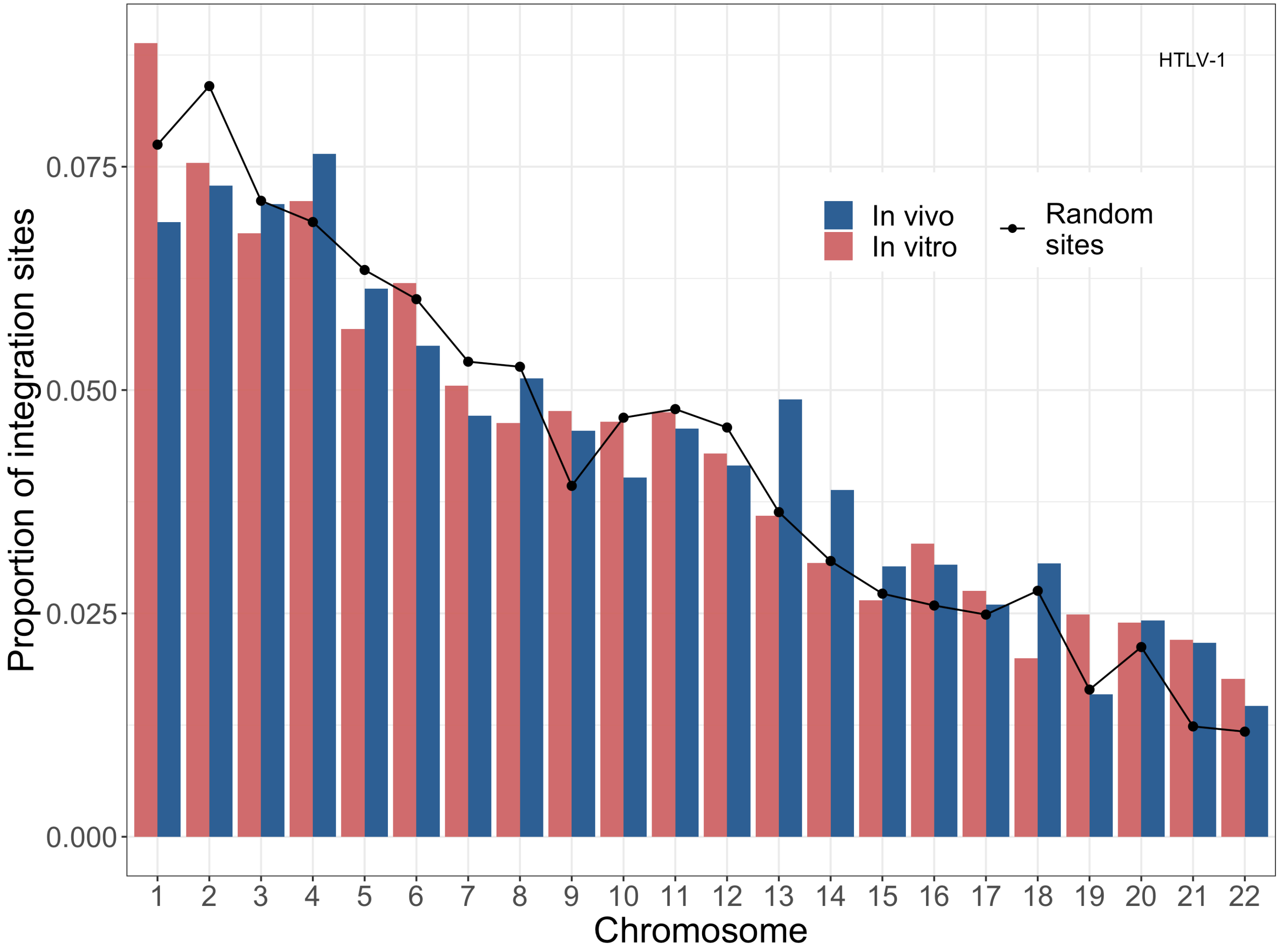


***Fig. S1. HTLV-1 Integration site survival is biased to specific chromosomes.*** *For each chromosome, the relative frequency of integration sites present in vivo and in vitro is shown (in vivo - red, in vitro - blue). The black line shows the proportion of random sites found in each chromosome.*

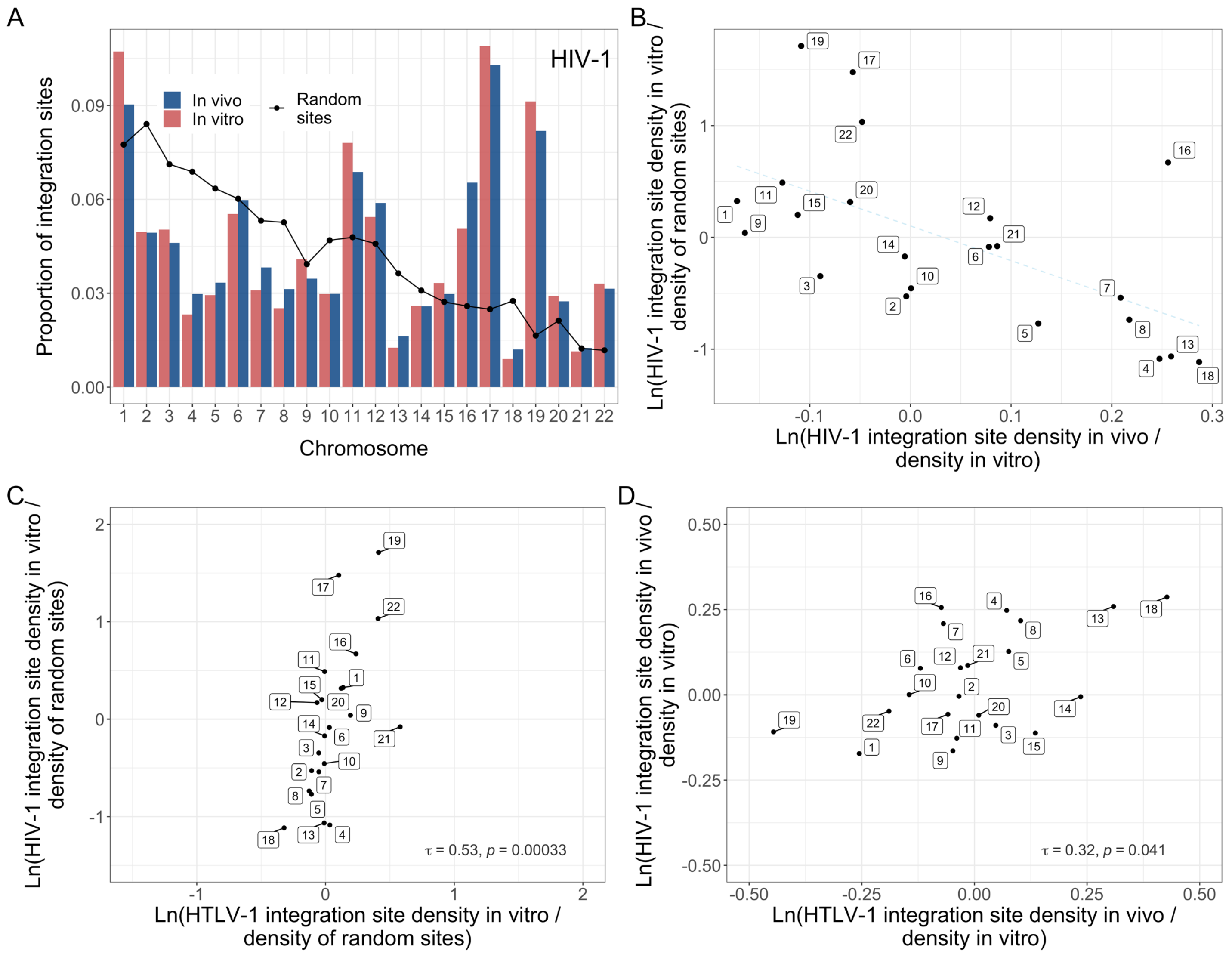


***Fig. S2. HIV-1 integration site targeting and survival is biased to specific chromosomes.*** *For each chromosome, the relative frequency of integration sites present in the HIV-1 in vitro and HIV-1 in vivo datasets was calculated.* ***(A)*** *Integration site frequency. The black line shows the proportion of random sites found in each chromosome.* ***(B)*** *For each chromosome, two ratios of integration site frequencies (F) were calculated:* $F_{in\ vivo}\ /\ F_{in\ vitro}$ *and* $F_{in\ vitro}\ /\ F_{random\ sites}$ *(logarithmic scales). As in HTLV-1 (Fig. 1), HIV-1 survival is most strongly favoured in chromosome 18.* ***(C)*** *Significant positive correlation between the rank order of chromosomes favoured for initial integration between HTLV-1 and HIV-1 (Kendall's rank correlation test); the magnitude of chromosome preference in initial integration targeting is much greater in HIV-1.* ***(D)*** *Significant positive correlation between HTLV-1 and HIV-1 in the rank order of chromosomes favoured for integration site survival in vivo (Kendall's rank correlation test).*

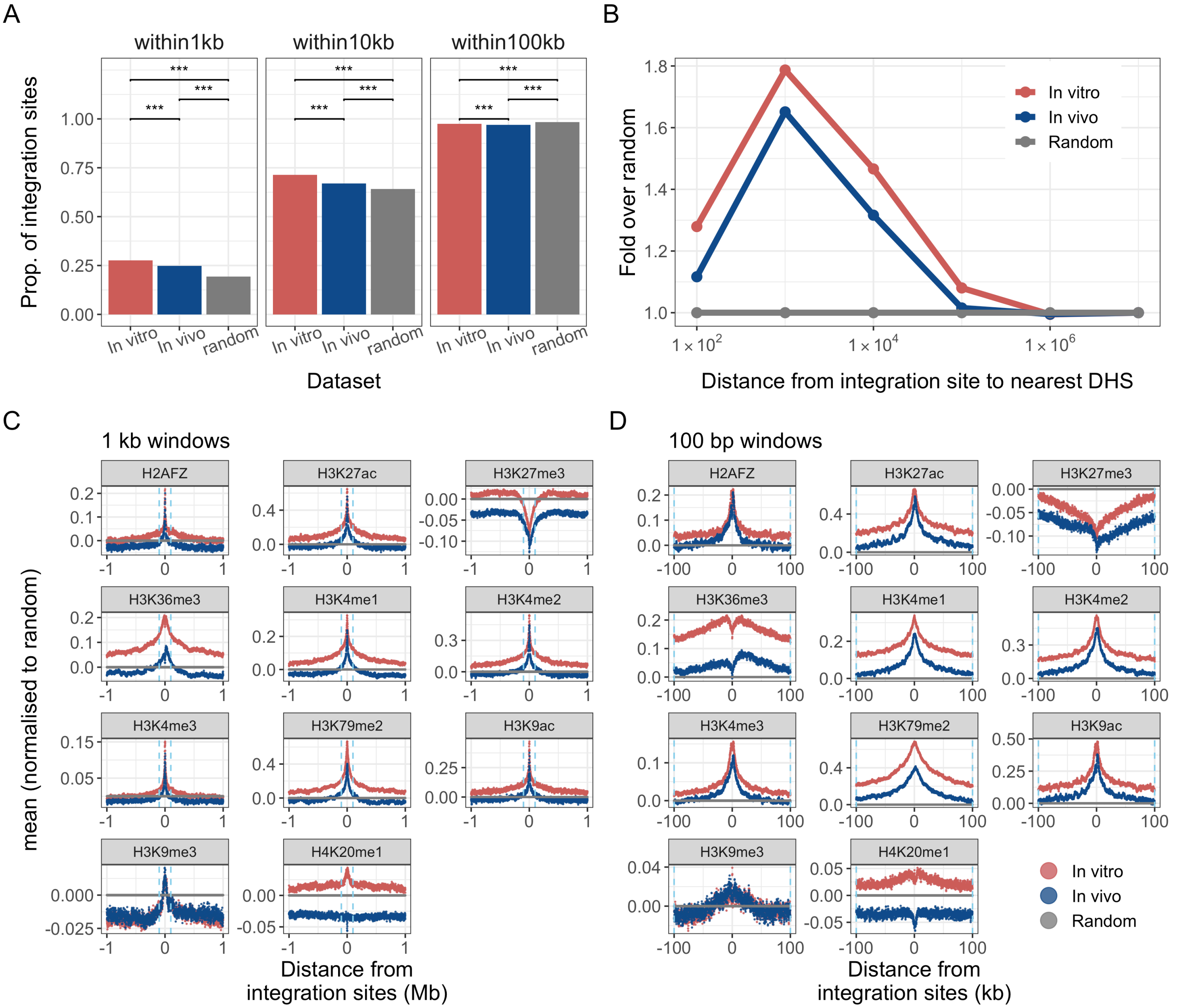


***Fig. S3. Initial HTLV-1 integration is biased towards accessible, transcriptionally active chromatin.*** *HTLV-1 Integration site and random sites were mapped with respect to transcription factor binding sites (TFBS), DNAse hypersensitive sites (DHS) and histone marks mapped by the ENCODE project for the B cell line GM12878. (A) In vitro sites were annotated with respect to the distance to the nearest binding site of any of the transcription factors . HTLV-1 integration sites both in vitro and in vivo were significantly enriched within 1 kb and 10 kb of any TFBS (***   $p < 10^{-16}$; $\chi^2$ test with Bonferroni correction). (B) Integration sites were enriched within 10 kb of a DNAse hypersensitive site (DHS). In each analysis, integration site frequency is compared to a random distribution. (C),(D) The mean density of each of 11 histone marks flanking the integration sites was calculated in discrete windows upstream and downstream across all integration sites from each dataset, either (C) - 1 kb windows, up to 1 Mb from the integration site, or (D) - 100bp windows, up to 100 kb from the integration site. $Ln(fold\ change\ from\ random)$ is shown. Dashed line denotes 100 kb upstream or downstream.*

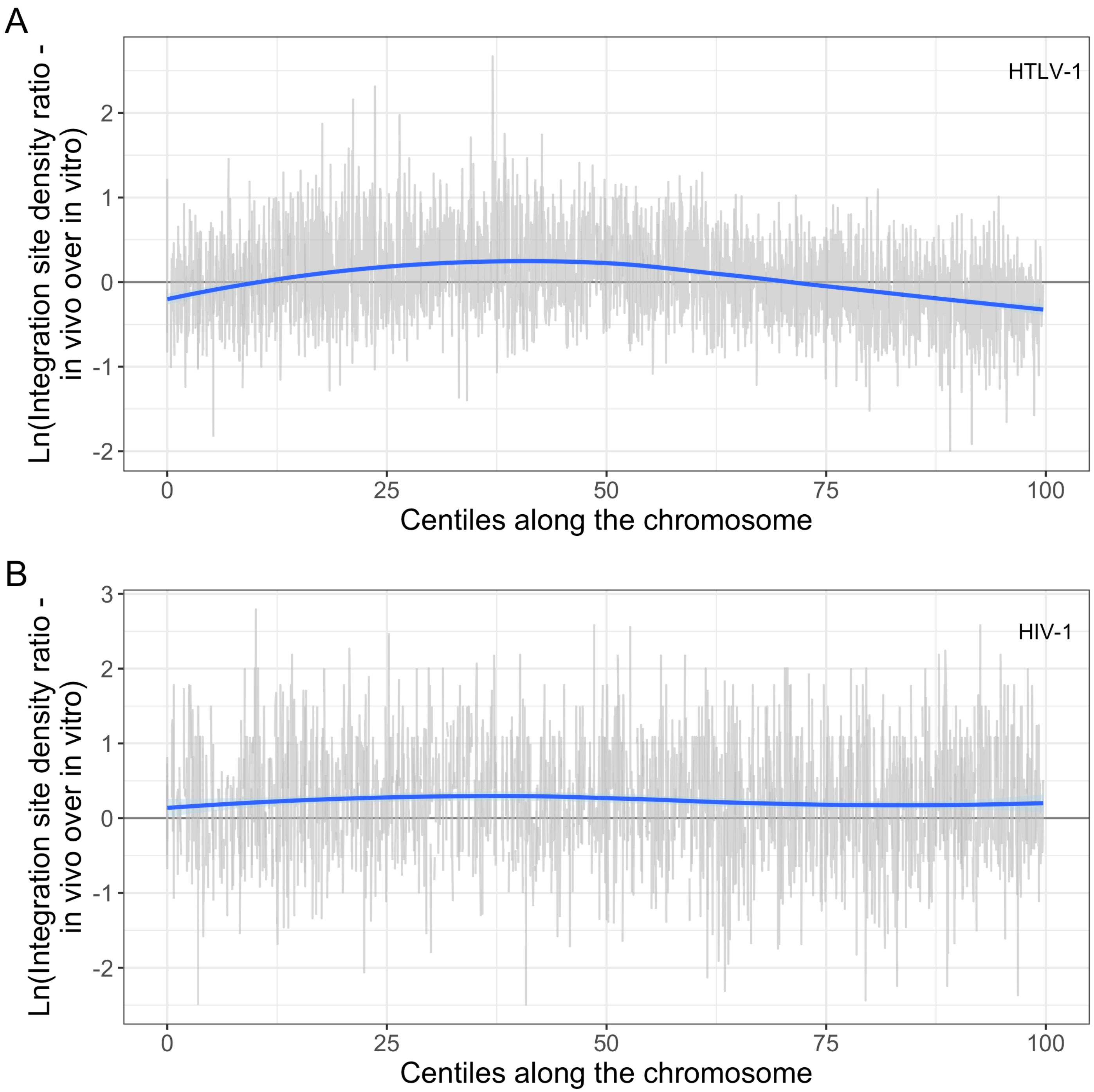


***Fig. S4. HTLV-1 and HIV-1 survival vs position along the chromosome.*** *For each chromosome, we calculated the clone survival index (CSI; of either HTLV-1* ***(A)*** *or HIV-1* ***(B)****) in discrete 1 Mb windows. Then for each chromosome, the genomic coordinates were converted to relative centiles along the chromosome, and the CSI plotted versus this relative position. In HTLV-1 in particular there is a clear trend (trendline using LOESS method) from increased survival in the centre of the chromosome to decreased survival towards the telomeres.*

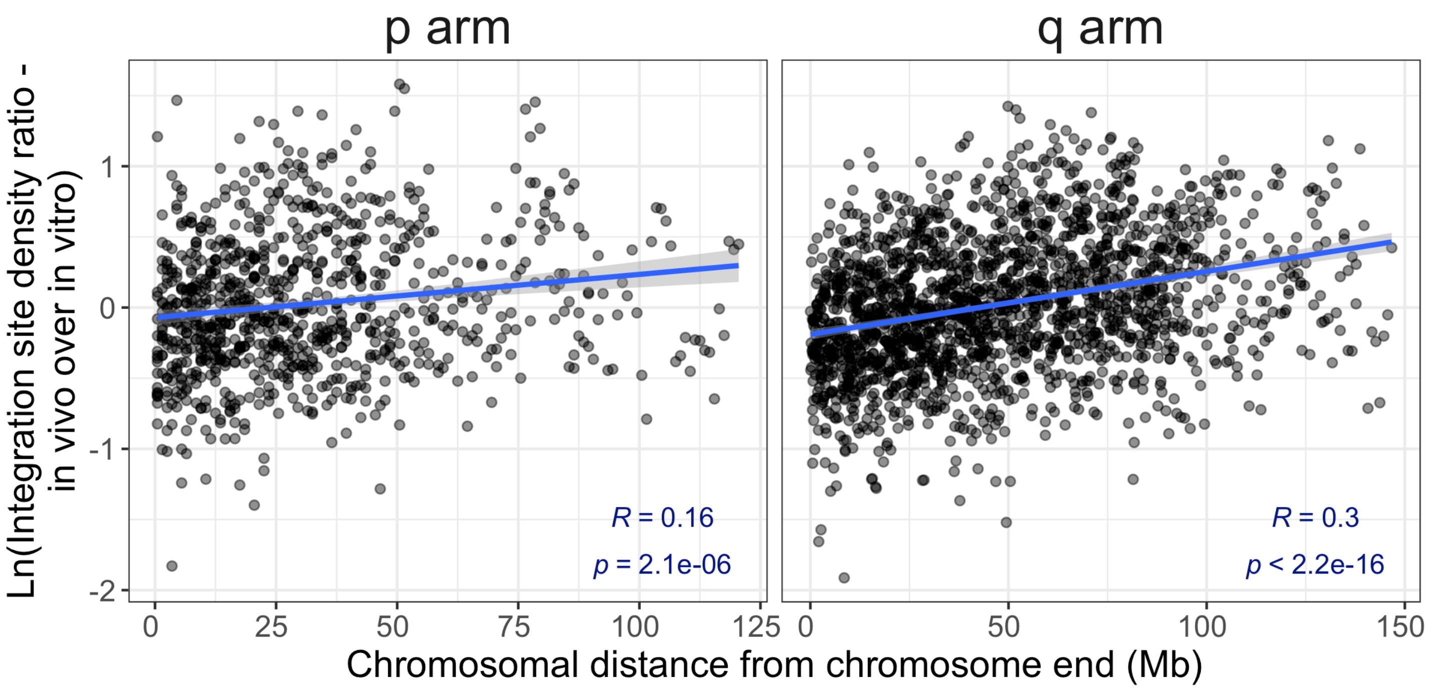


***Fig. S5. HTLV-1 survival vs distance from the telomere**. The HTLV-1 clone survival index $CSI_{HTLV-1}$ is significantly positively correlated with the absolute genomic distance from the telomere on both short and long arms of the chromosome (Pearson's correlation test); however, this correlation is weaker than the correlation with the distance from the centromere (Fig. 3).*

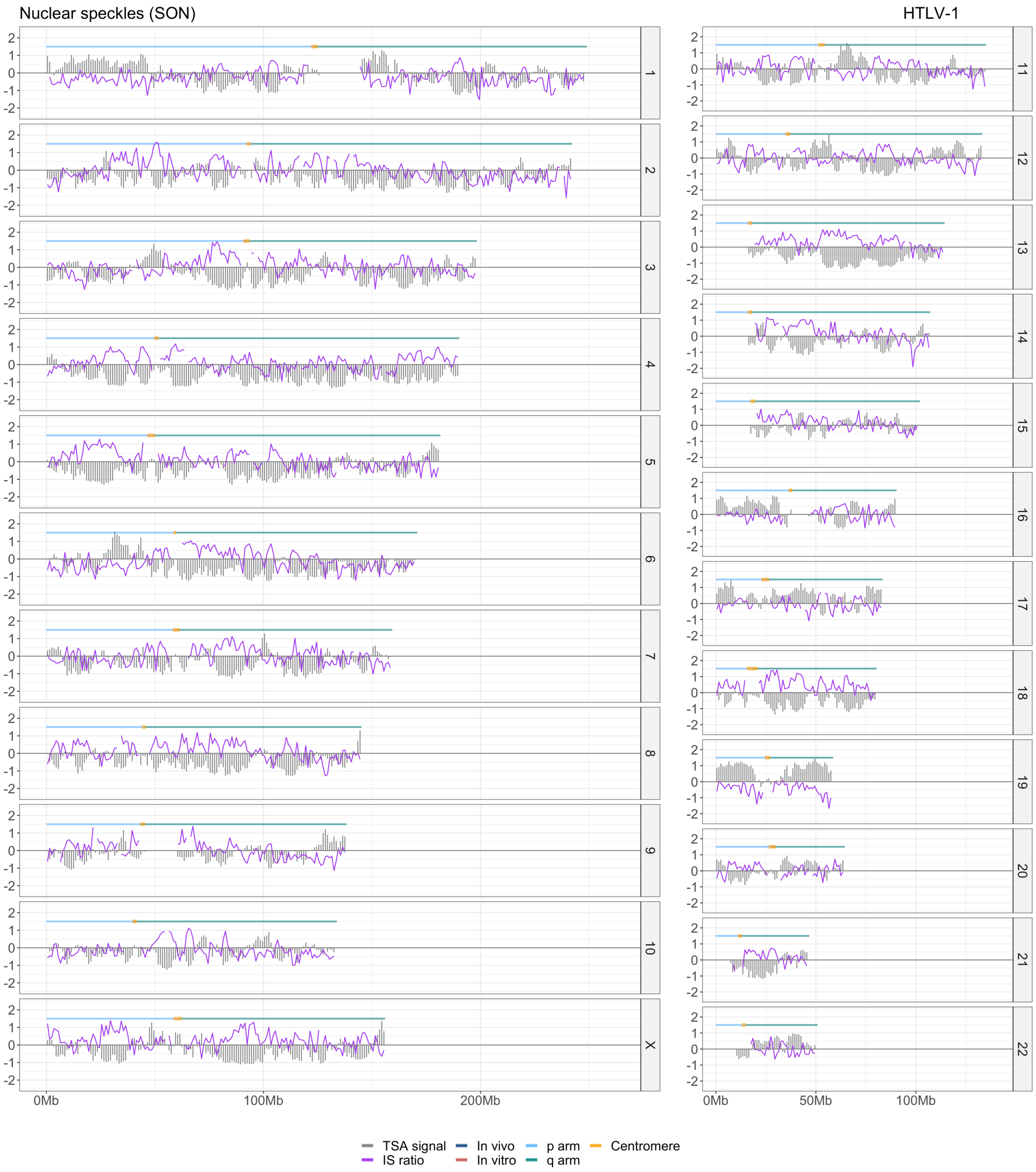


***Fig. S6. HTLV-1 Clone survival index vs distance from nuclear speckles.*** *In each panel, the TSA-seq data for SON are plotted against* $CSI_{HTLV-1}$ *for each chromosome. The panels for chromosomes 11, 12 are also shown in Fig. 4B, and are included here for completeness. The panel for chromosome 10 is also included in Fig. 5A.*

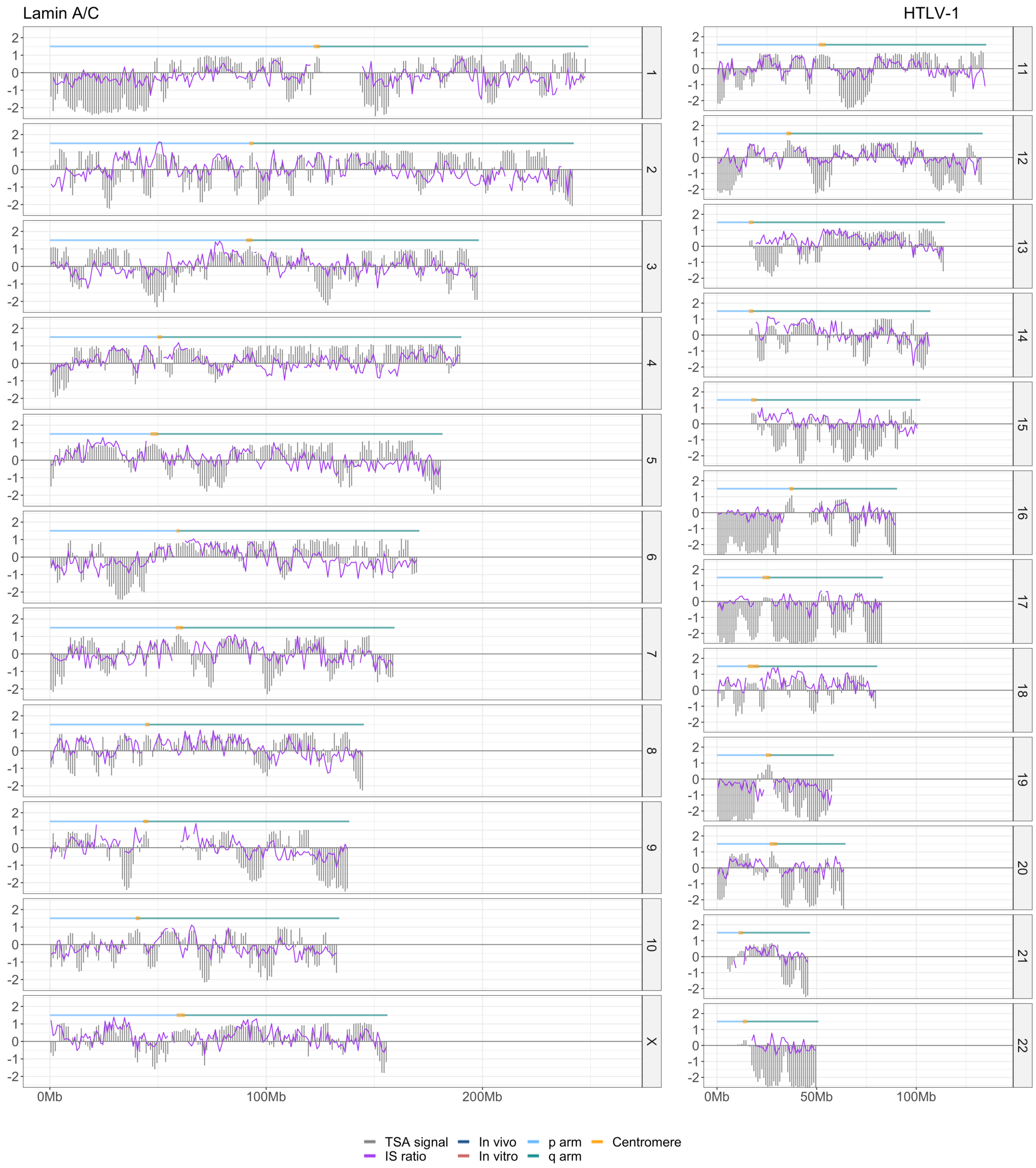


***Fig. S7. HTLV-1 Clone survival index vs distance from Lamina**. In each panel, the TSA-seq data for Lamin A/C are plotted against $CSI_{HTLV-1}$ for each chromosome. The panels for chromosomes 11, 12 are also included in Fig. 4B. The panel for chromosome 10 is also included in Fig. 5A.*

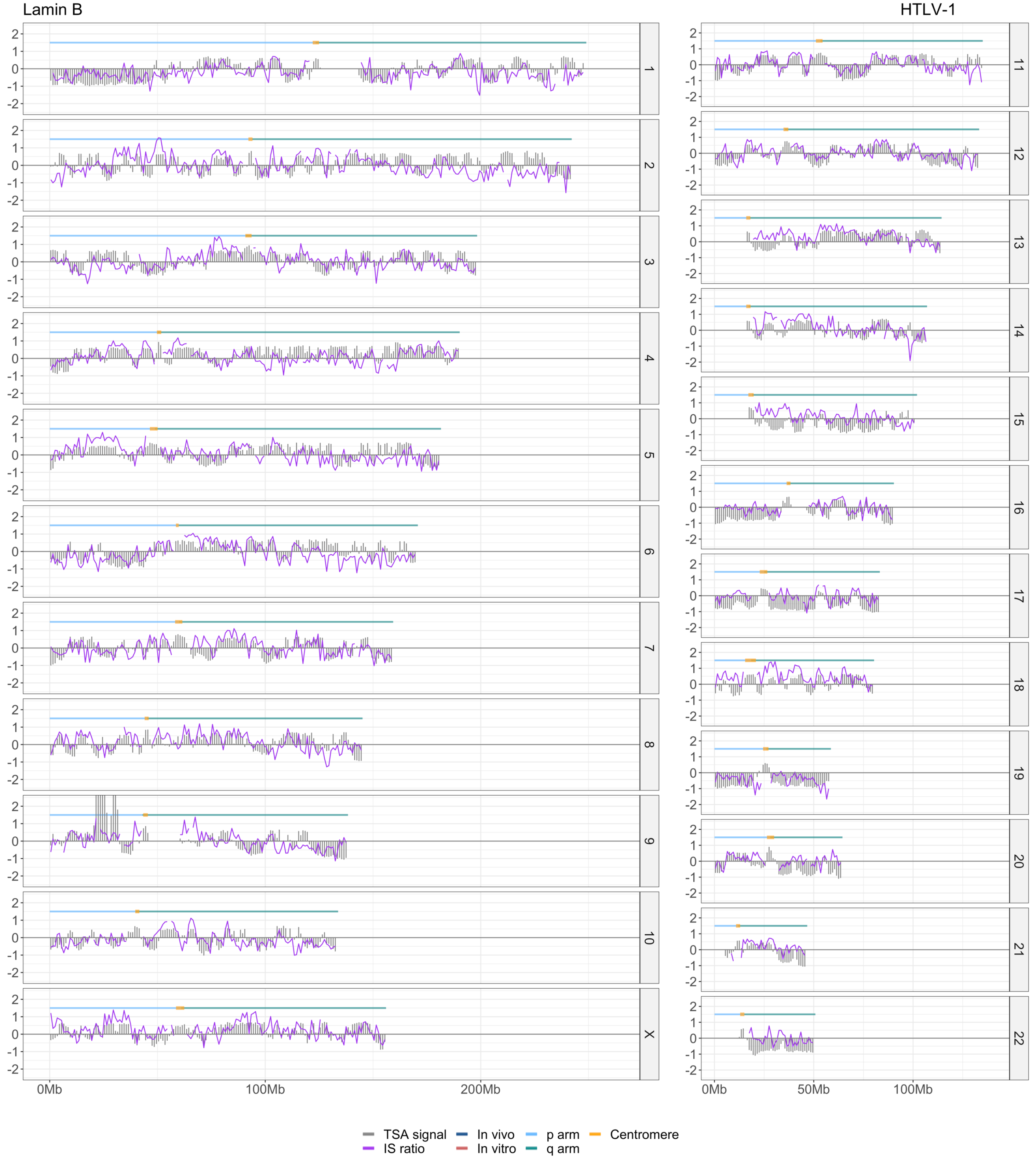


***Fig. S8. HTLV-1 Clone survival index vs distance from Lamina.*** *In each panel, the TSA-seq data for Lamin B are plotted against* $CSI_{HTLV-1}$ *for each chromosome.*

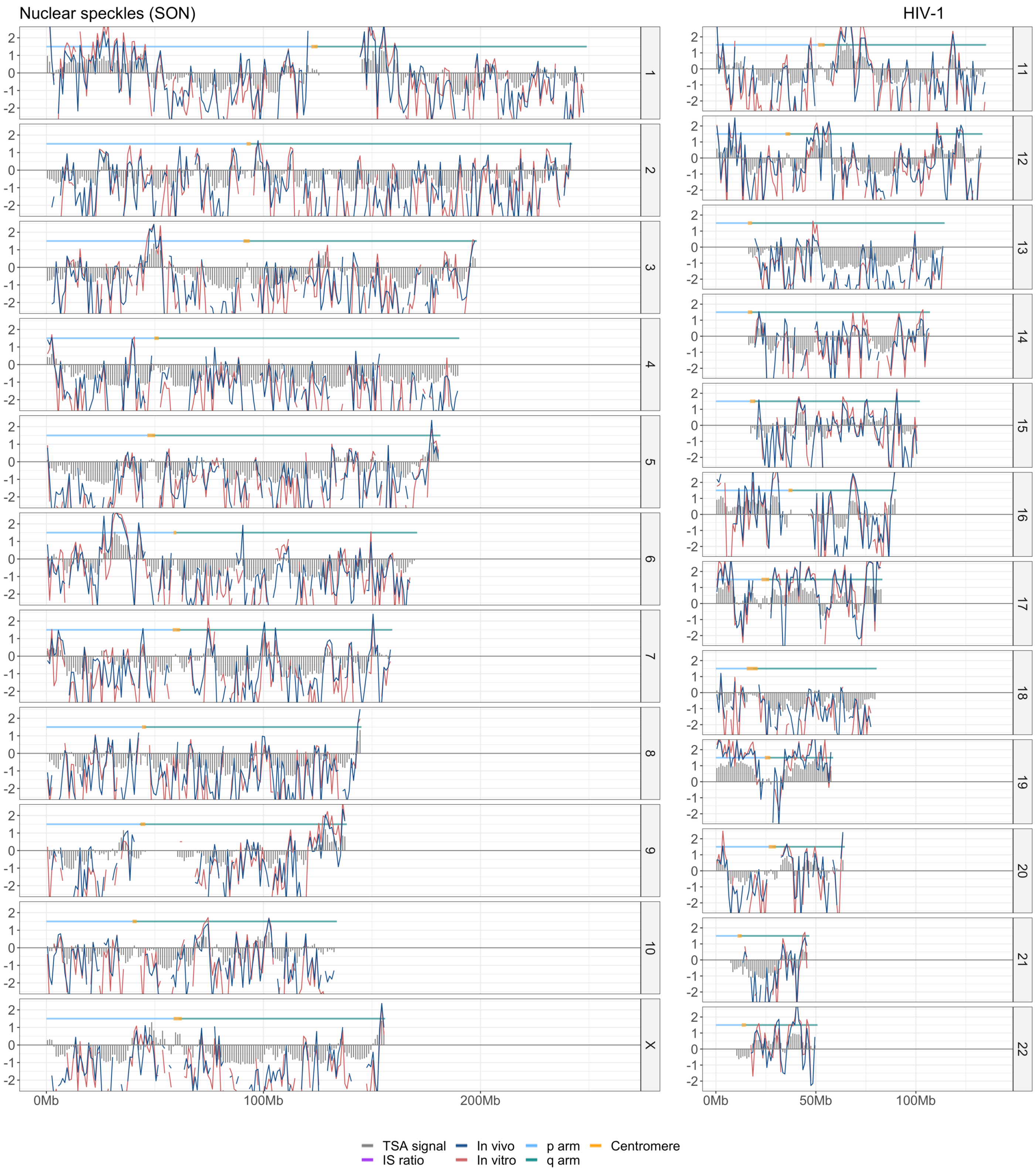


***Fig. S9. HIV-1 integration site frequency vs distance from nuclear speckles.*** *In each panel, the TSA-seq data for SON are plotted against the integration site frequency for either HIV-1 in vitro (red) or in vivo (blue) (normalized over random) in each chromosome.*

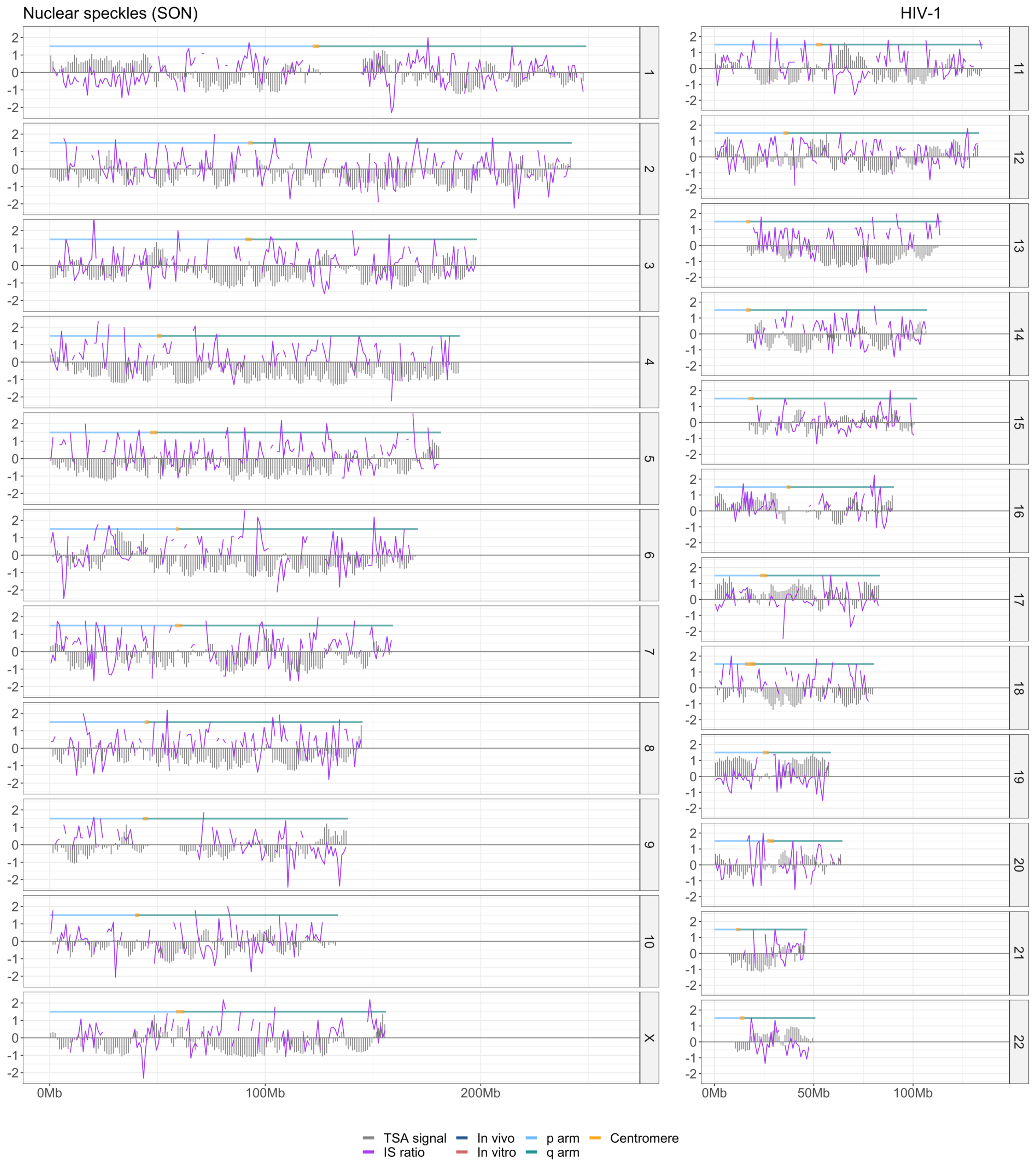


***Fig. S10. HIV-1 Clone survival index vs distance from nuclear speckles.*** *In each panel, the TSA-seq data for SON are plotted against* $CSI_{HIV-1}$ *for each chromosome. The panels for chromosomes 11, 12 are also shown in Fig. 4C, and included here for completeness.*

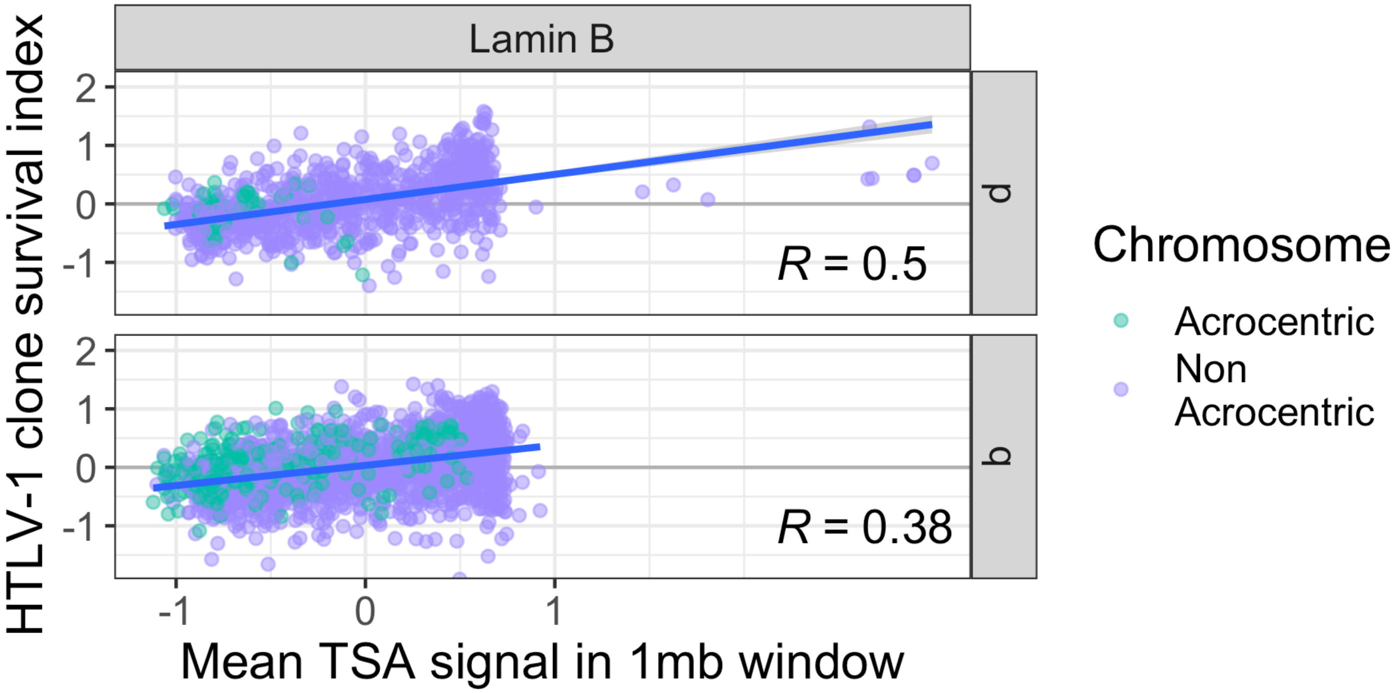


***Fig. S11. Genome-wide correlation between selective HTLV-1 proviral survival and proximity to nuclear lamina.*** *TSA-seq data on Lamin-B are significantly correlated with the clone survival index across the whole genome (Pearson's correlation test, $p < 10^{-16}$ for each correlation).*

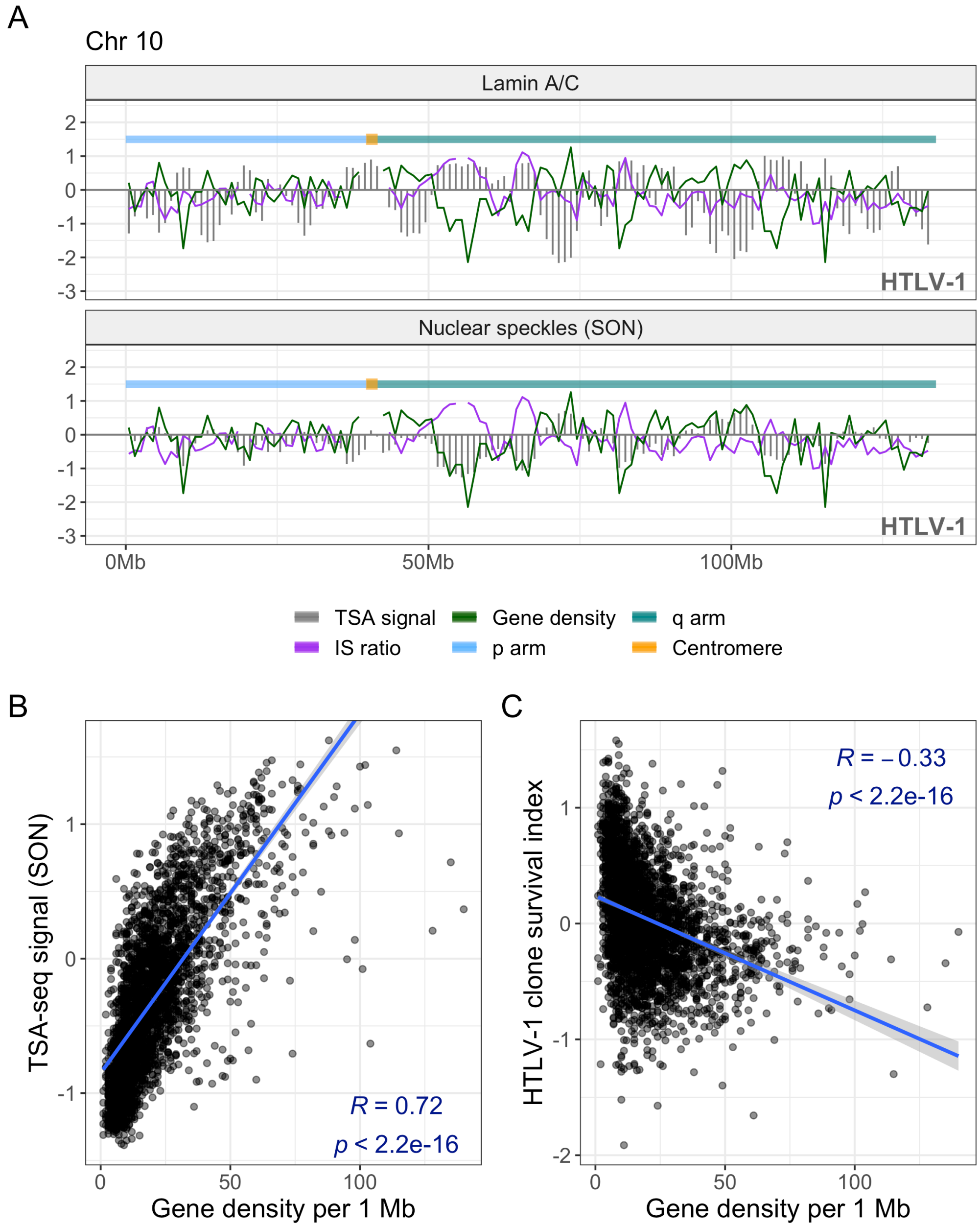


***Fig. S12. HTLV-1 clone survival is reduced in gene-rich genomic regions.*** *The number of genes overlapping each 1 Mb window across each chromosome was quantified and compared against the HTLV-1 clone survival index.* ***(A)*** *TSA-seq data on Lamin A/C and SON from (20) are plotted against the gene density (green line) and the HTLV-1 clone survival index* $CSI_{HTLV-1}$ *(purple). The gene density closely follows the proximity to nuclear speckles.* ***(B)*** *Genome-wide, there is a significant positive correlation (Pearson's correlation test) between the proximity to nuclear speckles and the gene density.* ***(C)*** *A significant negative correlation is observed between the HTLV-1 clone survival index and gene density.*

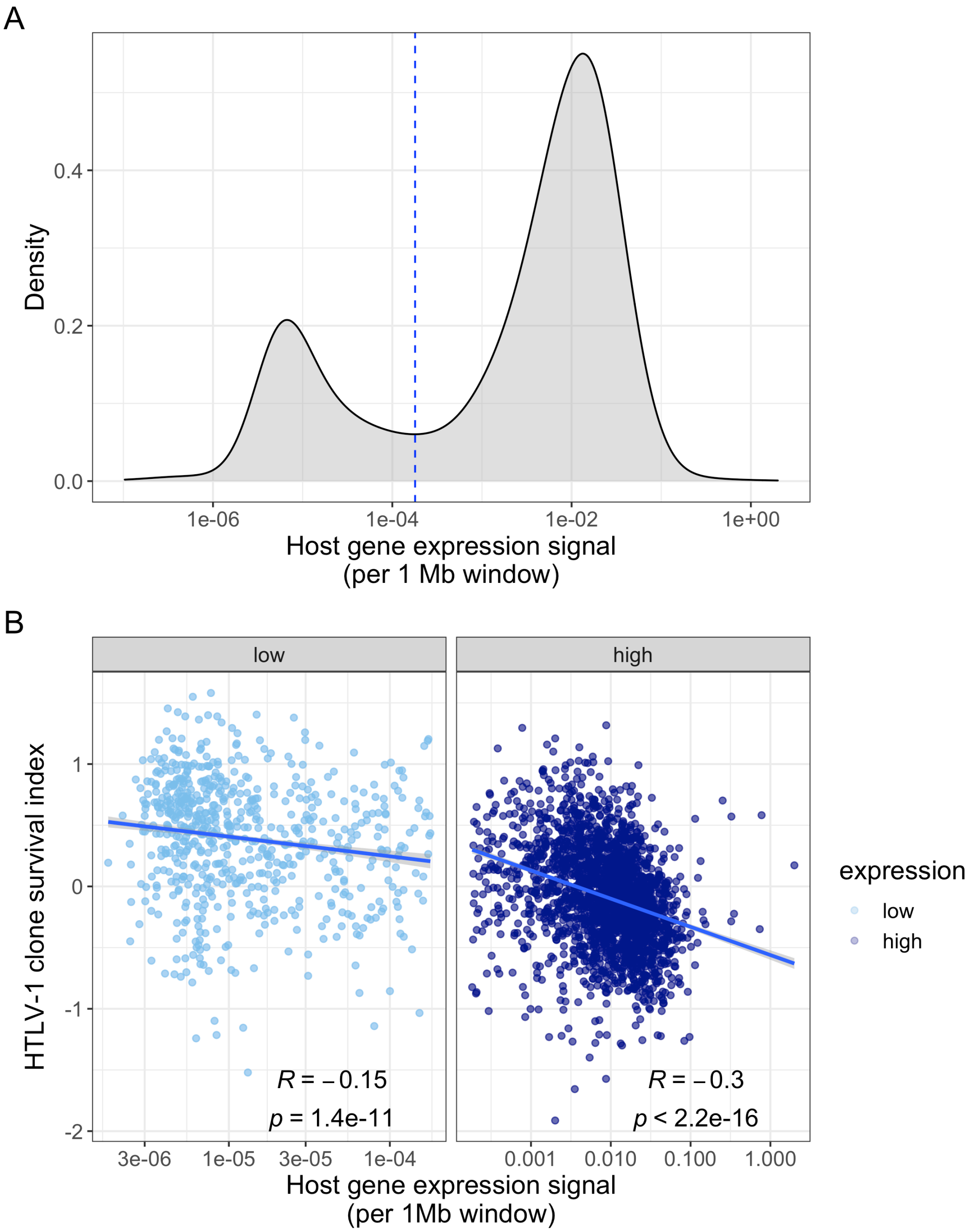


***Fig. S13. Counterselection of HTLV-1 proviruses in highly expressing genomic regions****. The average RNA expression intensity per 1 Mb window was quantified across each chromosome.* ***(A)*** *the distribution of RNA expression density (shown on log scale) is bimodal. A local minimum separates genomic regions with high expression from those with low expression.* ***(B)*** *The observed negative correlation between expression intensity and the* $CSI_{HTLV-1}$ *(Fig. 6) is also observed independently for low- and high-expressing regions (Pearson's correlation test).*

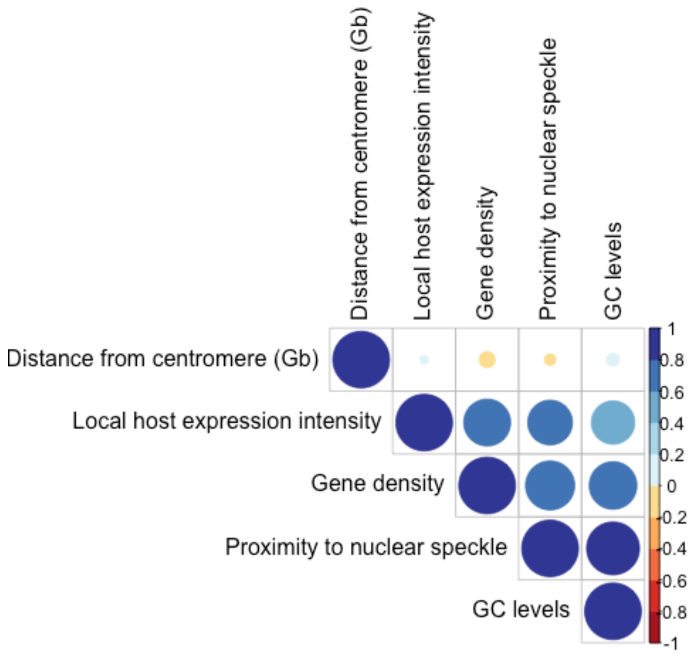


***Fig. S14. Correlation plot, potential predictors of HTLV-1 survival***

| | Dataset | Samples | Integration sites | Publication |
|---|---|---|---|---|
| HTLV-1 | HTLV1invivo1 | 142 | 225425 | (*1*) |
| | HTLV1invivo2 | 14 | 40410 | (*2*) |
| | HTLV1invivo3 | 196 | 6393 | (*3*) |
| | HTLV1invivo4 | 36 | 60350 | (*4*) |
| | HTLV1invitro1 | 5 | 4687 | (*1*) |
| | HTLV1invitro2 | 3 | 4444 | (*2*) |
| | HTLV1invitro3 | 8 | 226019 | This manuscript |
| HIV-1 | HIV1invivo1 | 36 | 32569 | (*5*) |
| | HIV1invivo2 | 25 | 13142 | (*5*) |
| | HIV1invitro1 | 2 | 65924 | (*5*) |
| Random | In silico sites | 1 | 109218 | This manuscript |

**Table S1** HTLV-1 integration site datasets used in this work

| Dataset type | file accession | experiment/ project accession | assay name | target | lab | data availability |
|---|---|---|---|---|---|---|
| TFBS | ENCFF003VDB | ENCSR778UBR | ChIP-seq | ARID3A | Michael Snyder, Stanford | www.encodeproject.org |
| TFBS | ENCFF758RQJ | ENCSR590KEQ | ChIP-seq | ARNT | Michael Snyder, Stanford | www.encodeproject.org |
| TFBS | ENCFF096XRG | ENCSR849WCQ | ChIP-seq | ASH2L | Bradley Bernstein, Broad | www.encodeproject.org |
| TFBS | ENCFF806KKM | ENCSR000BQK | ChIP-seq | ATF2 | Richard Myers, HAIB | www.encodeproject.org |
| TFBS | ENCFF495PWL | ENCSR014YCR | ChIP-seq | ATF7 | Michael Snyder, Stanford | www.encodeproject.org |
| TFBS | ENCFF725YZH | ENCSR636MKU | ChIP-seq | BACH1 | Michael Snyder, Stanford | www.encodeproject.org |
| TFBS | ENCFF832YIE | ENCSR000BGT | ChIP-seq | BATF | Richard Myers, HAIB | www.encodeproject.org |
| TFBS | ENCFF383HAY | ENCSR000BHA | ChIP-seq | BCL11A | Richard Myers, HAIB | www.encodeproject.org |
| TFBS | ENCFF247MHT | ENCSR000BNQ | ChIP-seq | BCL3 | Richard Myers, HAIB | www.encodeproject.org |
| TFBS | ENCFF587BJK | ENCSR000BJZ | ChIP-seq | BCLAF1 | Richard Myers, HAIB | www.encodeproject.org |
| TFBS | ENCFF370ZNL | ENCSR987MTA | ChIP-seq | BHLHE40 | Michael Snyder, Stanford | www.encodeproject.org |
| TFBS | ENCFF592LPO | ENCSR469WII | ChIP-seq | BMI1 | Michael Snyder, Stanford | www.encodeproject.org |
| TFBS | ENCFF005JKU | ENCSR000DZS | ChIP-seq | BRCA1 | Michael Snyder, Stanford | www.encodeproject.org |
| TFBS | ENCFF070SOX | ENCSR860UHK | ChIP-seq | CBFB | Richard Myers, HAIB | www.encodeproject.org |
| TFBS | ENCFF552QOA | ENCSR549NPZ | ChIP-seq | CBX3 | Bradley Bernstein, Broad | www.encodeproject.org |
| TFBS | ENCFF417SVR | ENCSR372GIN | ChIP-seq | CBX5 | Richard Myers, HAIB | www.encodeproject.org |
| TFBS | ENCFF786YYI | ENCSR681NOM | ChIP-seq | CEBPB | Michael Snyder, Stanford | www.encodeproject.org |
| TFBS | ENCFF243GOG | ENCSR347NOB | ChIP-seq | CEBPZ | Michael Snyder, Stanford | www.encodeproject.org |
| TFBS | ENCFF863CTN | ENCSR000DZE | ChIP-seq | CHD1 | Michael Snyder, Stanford | www.encodeproject.org |
| TFBS | ENCFF546AYN | ENCSR000DZR | ChIP-seq | CHD2 | Michael Snyder, Stanford | www.encodeproject.org |
| TFBS | ENCFF249SIN | ENCSR751CJG | ChIP-seq | CHD4 | Bradley Bernstein, Broad | www.encodeproject.org |
| TFBS | ENCFF091YID | ENCSR839XZU | ChIP-seq | CREM | Richard Myers, HAIB | www.encodeproject.org |
| TFBS | ENCFF960ZGP | ENCSR000DZN | ChIP- | CTCF | Michael Snyder, | www.encodeproject.org |

| Dataset type | file accession | experiment/ project accession | assay name | target | lab | data availability |
|---|---|---|---|---|---|---|
| | | | seq | | Stanford | |
| TFBS | ENCFF567NFS | ENCSR000DYR | ChIP-seq | CUX1 | Michael Snyder, Stanford | www.encodeproject.org |
| TFBS | ENCFF771IAW | ENCSR509FWH | ChIP-seq | DPF2 | Michael Snyder, Stanford | www.encodeproject.org |
| TFBS | ENCFF687SFB | ENCSR000DYY | ChIP-seq | E2F4 | Michael Snyder, Stanford | www.encodeproject.org |
| TFBS | ENCFF412GFI | ENCSR793HVL | ChIP-seq | E2F8 | Michael Snyder, Stanford | www.encodeproject.org |
| TFBS | ENCFF035GFS | ENCSR439WAF | ChIP-seq | E4F1 | Michael Snyder, Stanford | www.encodeproject.org |
| TFBS | ENCFF864NSW | ENCSR000BGU | ChIP-seq | EBF1 | Richard Myers, HAIB | www.encodeproject.org |
| TFBS | ENCFF023ALY | ENCSR199WXF | ChIP-seq | EED | Richard Myers, HAIB | www.encodeproject.org |
| TFBS | ENCFF948CPI | ENCSR841NDX | ChIP-seq | ELF1 | Michael Snyder, Stanford | www.encodeproject.org |
| TFBS | ENCFF432AQP | ENCSR000DZB | ChIP-seq | ELK1 | Michael Snyder, Stanford | www.encodeproject.org |
| TFBS | ENCFF080HJX | ENCSR000DZG | ChIP-seq | EP300 | Michael Snyder, Stanford | www.encodeproject.org |
| TFBS | ENCFF722LJP | ENCSR000DYQ | ChIP-seq | ESRRA | Michael Snyder, Stanford | www.encodeproject.org |
| TFBS | ENCFF980VOD | ENCSR000BKA | ChIP-seq | ETS1 | Richard Myers, HAIB | www.encodeproject.org |
| TFBS | ENCFF745ANU | ENCSR626VUC | ChIP-seq | ETV6 | Richard Myers, HAIB | www.encodeproject.org |
| TFBS | ENCFF615NYO | ENCSR000ARD | ChIP-seq | EZH2 | Bradley Bernstein, Broad | www.encodeproject.org |
| TFBS | ENCFF990MTR | ENCSR861JUQ | ChIP-seq | FOXK2 | Michael Snyder, Stanford | www.encodeproject.org |
| TFBS | ENCFF946ACA | ENCSR331HPA | ChIP-seq | GABPA | Richard Myers, HAIB | www.encodeproject.org |
| TFBS | ENCFF298AIX | ENCSR828NCB | ChIP-seq | GATAD2B | Michael Snyder, Stanford | www.encodeproject.org |
| TFBS | ENCFF722QBB | ENCSR514VAY | ChIP-seq | HCFC1 | Michael Snyder, Stanford | www.encodeproject.org |
| TFBS | ENCFF299UPZ | ENCSR330OEO | ChIP-seq | HDAC2 | Bradley Bernstein, Broad | www.encodeproject.org |
| TFBS | ENCFF442WRJ | ENCSR145XQO | ChIP-seq | HDGF | Michael Snyder, Stanford | www.encodeproject.org |
| TFBS | ENCFF603BID | ENCSR009MBP | ChIP-seq | HSF1 | Michael Snyder, Stanford | www.encodeproject.org |
| TFBS | ENCFF018NNF | ENCSR441VHN | ChIP-seq | IKZF1 | Michael Snyder, Stanford | www.encodeproject.org |

| Dataset type | file accession | experiment/ project accession | assay name | target | lab | data availability |
|---|---|---|---|---|---|---|
| TFBS | ENCFF088OLI | ENCSR822AHX | ChIP-seq | IKZF2 | Michael Snyder, Stanford | www.encodeproject.org |
| TFBS | ENCFF604AZX | ENCSR408JQO | ChIP-seq | IRF3 | Michael Snyder, Stanford | www.encodeproject.org |
| TFBS | ENCFF720YMW | ENCSR000BGY | ChIP-seq | IRF4 | Richard Myers, HAIB | www.encodeproject.org |
| TFBS | ENCFF843HDK | ENCSR976TBC | ChIP-seq | IRF5 | Michael Snyder, Stanford | www.encodeproject.org |
| TFBS | ENCFF478XNA | ENCSR897MMC | ChIP-seq | JUNB | Richard Myers, HAIB | www.encodeproject.org |
| TFBS | ENCFF873DJD | ENCSR000DYS | ChIP-seq | JUND | Michael Snyder, Stanford | www.encodeproject.org |
| TFBS | ENCFF710ROZ | ENCSR000DNO | ChIP-seq | KAT2A | Kevin Struhl, HMS | www.encodeproject.org |
| TFBS | ENCFF799KZP | ENCSR391IWM | ChIP-seq | KDM1A | Bradley Bernstein, Broad | www.encodeproject.org |
| TFBS | ENCFF417WPC | ENCSR974OFJ | ChIP-seq | KLF5 | Michael Snyder, Stanford | www.encodeproject.org |
| TFBS | ENCFF305SLO | ENCSR657PEW | ChIP-seq | LARP7 | Michael Snyder, Stanford | www.encodeproject.org |
| TFBS | ENCFF186AWV | ENCSR000DYV | ChIP-seq | MAFK | Michael Snyder, Stanford | www.encodeproject.org |
| TFBS | ENCFF270NAL | ENCSR000DZF | ChIP-seq | MAX | Michael Snyder, Stanford | www.encodeproject.org |
| TFBS | ENCFF348STZ | ENCSR000DZA | ChIP-seq | MAZ | Michael Snyder, Stanford | www.encodeproject.org |
| TFBS | ENCFF958GXF | ENCSR000BKB | ChIP-seq | MEF2A | Richard Myers, HAIB | www.encodeproject.org |
| TFBS | ENCFF623FAW | ENCSR177VFS | ChIP-seq | MEF2B | Michael Snyder, Stanford | www.encodeproject.org |
| TFBS | ENCFF830BRO | ENCSR000BNG | ChIP-seq | MEF2C | Richard Myers, HAIB | www.encodeproject.org |
| TFBS | ENCFF125MEN | ENCSR552XSN | ChIP-seq | MLLT1 | Michael Snyder, Stanford | www.encodeproject.org |
| TFBS | ENCFF587POH | ENCSR293QAR | ChIP-seq | MTA2 | Michael Snyder, Stanford | www.encodeproject.org |
| TFBS | ENCFF661FMB | ENCSR000BRH | ChIP-seq | MTA3 | Richard Myers, HAIB | www.encodeproject.org |
| TFBS | ENCFF199HGX | ENCSR000DZI | ChIP-seq | MXI1 | Michael Snyder, Stanford | www.encodeproject.org |
| TFBS | ENCFF402TSJ | ENCSR819ATC | ChIP-seq | MYB | Michael Snyder, Stanford | www.encodeproject.org |
| TFBS | ENCFF811VEN | ENCSR278SQL | ChIP-seq | NBN | Michael Snyder, Stanford | www.encodeproject.org |
| TFBS | ENCFF138ZBJ | ENCSR000BQL | ChIP- | NFATC1 | Richard Myers, | www.encodeproject.org |

| Dataset type | file accession | experiment/ project accession | assay name | target | lab | data availability |
|---|---|---|---|---|---|---|
| | | | seq | | HAIB | |
| TFBS | ENCFF704PDA | ENCSR437GBJ | ChIP-seq | NFATC3 | Michael Snyder, Stanford | www.encodeproject.org |
| TFBS | ENCFF480WDX | ENCSR000BRN | ChIP-seq | NFIC | Richard Myers, HAIB | www.encodeproject.org |
| TFBS | ENCFF860IXB | ENCSR746XEG | ChIP-seq | NFXL1 | Michael Snyder, Stanford | www.encodeproject.org |
| TFBS | ENCFF278GJK | ENCSR000DNN | ChIP-seq | NFYA | Kevin Struhl, HMS | www.encodeproject.org |
| TFBS | ENCFF510NDO | ENCSR000DNM | ChIP-seq | NFYB | Kevin Struhl, HMS | www.encodeproject.org |
| TFBS | ENCFF084NXU | ENCSR732PJX | ChIP-seq | NKRF | Michael Snyder, Stanford | www.encodeproject.org |
| TFBS | ENCFF462AKP | ENCSR784VIQ | ChIP-seq | NR2C1 | Michael Snyder, Stanford | www.encodeproject.org |
| TFBS | ENCFF434HVY | ENCSR000EUL | ChIP-seq | NR2C2 | Peggy Farnham, USC | www.encodeproject.org |
| TFBS | ENCFF531KOV | ENCSR514VYD | ChIP-seq | NR2F1 | Michael Snyder, Stanford | www.encodeproject.org |
| TFBS | ENCFF652BRY | ENCSR000DZO | ChIP-seq | NRF1 | Michael Snyder, Stanford | www.encodeproject.org |
| TFBS | ENCFF946SAG | ENCSR000BHJ | ChIP-seq | PAX5 | Richard Myers, HAIB | www.encodeproject.org |
| TFBS | ENCFF992JWY | ENCSR192AFN | ChIP-seq | PAX8 | Michael Snyder, Stanford | www.encodeproject.org |
| TFBS | ENCFF926LHG | ENCSR000BGR | ChIP-seq | PBX3 | Richard Myers, HAIB | www.encodeproject.org |
| TFBS | ENCFF335ADU | ENCSR711XNY | ChIP-seq | PKNOX1 | Michael Snyder, Stanford | www.encodeproject.org |
| TFBS | ENCFF455ZLJ | ENCSR000BGD | ChIP-seq | POLR2A | Richard Myers, HAIB | www.encodeproject.org |
| TFBS | ENCFF847DXY | ENCSR000DZK | ChIP-seq | POLR2AphosphoS 2 | Michael Snyder, Stanford | www.encodeproject.org |
| TFBS | ENCFF600GQL | ENCSR000BIF | ChIP-seq | POLR2AphosphoS 5 | Richard Myers, HAIB | www.encodeproject.org |
| TFBS | ENCFF654EGO | ENCSR000BMY | ChIP-seq | RAD21 | Richard Myers, HAIB | www.encodeproject.org |
| TFBS | ENCFF996NBR | ENCSR482TWQ | ChIP-seq | RAD51 | Michael Snyder, Stanford | www.encodeproject.org |
| TFBS | ENCFF034OSV | ENCSR785OKZ | ChIP-seq | RB1 | Michael Snyder, Stanford | www.encodeproject.org |
| TFBS | ENCFF687SSY | ENCSR330EXS | ChIP-seq | RBBP5 | Bradley Bernstein, Broad | www.encodeproject.org |
| TFBS | ENCFF470ZMK | ENCSR000DZC | ChIP-seq | RCOR1 | Michael Snyder, Stanford | www.encodeproject.org |

| Dataset type | file accession | experiment/ project accession | assay name | target | lab | data availability |
|---|---|---|---|---|---|---|
| TFBS | ENCFF105YDI | ENCSR387QUV | ChIP-seq | RELB | Michael Snyder, Stanford | www.encodeproject.org |
| TFBS | ENCFF313CII | ENCSR000BQS | ChIP-seq | REST | Richard Myers, HAIB | www.encodeproject.org |
| TFBS | ENCFF259LNG | ENCSR000DZW | ChIP-seq | RFX5 | Michael Snyder, Stanford | www.encodeproject.org |
| TFBS | ENCFF677QUK | ENCSR000BRI | ChIP-seq | RUNX3 | Richard Myers, HAIB | www.encodeproject.org |
| TFBS | ENCFF313BDA | ENCSR000BJD | ChIP-seq | RXRA | Richard Myers, HAIB | www.encodeproject.org |
| TFBS | ENCFF050CYK | ENCSR000DYX | ChIP-seq | SIN3A | Michael Snyder, Stanford | www.encodeproject.org |
| TFBS | ENCFF864TFH | ENCSR000BJE | ChIP-seq | SIX5 | Richard Myers, HAIB | www.encodeproject.org |
| TFBS | ENCFF903KEI | ENCSR212YKD | ChIP-seq | SKIL | Michael Snyder, Stanford | www.encodeproject.org |
| TFBS | ENCFF987PGY | ENCSR813DCK | ChIP-seq | SMAD1 | Michael Snyder, Stanford | www.encodeproject.org |
| TFBS | ENCFF855SJG | ENCSR251OVJ | ChIP-seq | SMAD5 | Richard Myers, HAIB | www.encodeproject.org |
| TFBS | ENCFF052STI | ENCSR706YUH | ChIP-seq | SMARCA5 | Michael Snyder, Stanford | www.encodeproject.org |
| TFBS | ENCFF572RPI | ENCSR000DZP | ChIP-seq | SMC3 | Michael Snyder, Stanford | www.encodeproject.org |
| TFBS | ENCFF071ZMW | ENCSR000BGQ | ChIP-seq | SPI1 | Richard Myers, HAIB | www.encodeproject.org |
| TFBS | ENCFF766WWB | ENCSR041XML | ChIP-seq | SRF | Michael Snyder, Stanford | www.encodeproject.org |
| TFBS | ENCFF323QQU | ENCSR332EYT | ChIP-seq | STAT1 | Michael Snyder, Stanford | www.encodeproject.org |
| TFBS | ENCFF923CHO | ENCSR000DZV | ChIP-seq | STAT3 | Michael Snyder, Stanford | www.encodeproject.org |
| TFBS | ENCFF383YEA | ENCSR000BQZ | ChIP-seq | STAT5A | Richard Myers, HAIB | www.encodeproject.org |
| TFBS | ENCFF069YVD | ENCSR000DNP | ChIP-seq | SUPT20H | Kevin Struhl, HMS | www.encodeproject.org |
| TFBS | ENCFF540AAP | ENCSR000BGS | ChIP-seq | TAF1 | Richard Myers, HAIB | www.encodeproject.org |
| TFBS | ENCFF668JHK | ENCSR412QBS | ChIP-seq | TARDBP | Michael Snyder, Stanford | www.encodeproject.org |
| TFBS | ENCFF392JWA | ENCSR000DYZ | ChIP-seq | TBL1XR1 | Michael Snyder, Stanford | www.encodeproject.org |
| TFBS | ENCFF896UZB | ENCSR000DZZ | ChIP-seq | TBP | Michael Snyder, Stanford | www.encodeproject.org |
| TFBS | ENCFF971VHK | ENCSR739IHN | ChIP- | TBX21 | Michael Snyder, | www.encodeproject.org |

| Dataset type | file accession | experiment/ project accession | assay name | target | lab | data availability |
|---|---|---|---|---|---|---|
| | | | seq | | Stanford | |
| TFBS | ENCFF768VSH | ENCSR000BGZ | ChIP-seq | TCF12 | Richard Myers, HAIB | www.encodeproject.org |
| TFBS | ENCFF152RNE | ENCSR501DKS | ChIP-seq | TCF7 | Richard Myers, HAIB | www.encodeproject.org |
| TFBS | ENCFF552WAH | ENCSR835XKS | ChIP-seq | TRIM22 | Michael Snyder, Stanford | www.encodeproject.org |
| TFBS | ENCFF295ZLM | ENCSR459FTB | ChIP-seq | UBTF | Michael Snyder, Stanford | www.encodeproject.org |
| TFBS | ENCFF514SWA | ENCSR000DZU | ChIP-seq | USF2 | Michael Snyder, Stanford | www.encodeproject.org |
| TFBS | ENCFF514DDI | ENCSR000EAA | ChIP-seq | WRNIP1 | Michael Snyder, Stanford | www.encodeproject.org |
| TFBS | ENCFF500RBO | ENCSR205SKQ | ChIP-seq | YBX1 | Michael Snyder, Stanford | www.encodeproject.org |
| TFBS | ENCFF223MUF | ENCSR000BNP | ChIP-seq | YY1 | Richard Myers, HAIB | www.encodeproject.org |
| TFBS | ENCFF630FLK | ENCSR207PFI | ChIP-seq | ZBED1 | Richard Myers, HAIB | www.encodeproject.org |
| TFBS | ENCFF475DID | ENCSR542FLV | ChIP-seq | ZBTB33 | Michael Snyder, Stanford | www.encodeproject.org |
| TFBS | ENCFF084IUW | ENCSR189YYK | ChIP-seq | ZBTB40 | Michael Snyder, Stanford | www.encodeproject.org |
| TFBS | ENCFF224WII | ENCSR900XDB | ChIP-seq | ZFP36 | Michael Snyder, Stanford | www.encodeproject.org |
| TFBS | ENCFF193POQ | ENCSR000DZL | ChIP-seq | ZNF143 | Michael Snyder, Stanford | www.encodeproject.org |
| TFBS | ENCFF676BIG | ENCSR117KWH | ChIP-seq | ZNF207 | Michael Snyder, Stanford | www.encodeproject.org |
| TFBS | ENCFF200SLC | ENCSR764CZW | ChIP-seq | ZNF217 | Michael Snyder, Stanford | www.encodeproject.org |
| TFBS | ENCFF313HBL | ENCSR072PWP | ChIP-seq | ZNF24 | Michael Snyder, Stanford | www.encodeproject.org |
| TFBS | ENCFF942MDT | ENCSR000DYP | ChIP-seq | ZNF384 | Michael Snyder, Stanford | www.encodeproject.org |
| TFBS | ENCFF615DTQ | ENCSR173ZVL | ChIP-seq | ZNF592 | Michael Snyder, Stanford | www.encodeproject.org |
| TFBS | ENCFF777DVJ | ENCSR075FNZ | ChIP-seq | ZNF622 | Michael Snyder, Stanford | www.encodeproject.org |
| TFBS | ENCFF137BRA | ENCSR859FDL | ChIP-seq | ZNF687 | Michael Snyder, Stanford | www.encodeproject.org |
| TFBS | ENCFF214NJL | ENCSR412YGM | ChIP-seq | ZSCAN29 | Michael Snyder, Stanford | www.encodeproject.org |
| TFBS | ENCFF260NAX | ENCSR000DNQ | ChIP-seq | ZZZ3 | Kevin Struhl, HMS | www.encodeproject.org |

| Dataset type | file accession | experiment/ project accession | assay name | target | lab | data availability |
|---|---|---|---|---|---|---|
| HistMarks | ENCFF831ZHL | ENCSR000AKF | ChIP-seq | H3K4me1 | Bradley Bernstein, Broad | www.encodeproject.org |
| HistMarks | ENCFF039HDL | ENCSR000AKG | ChIP-seq | H3K4me2 | Bradley Bernstein, Broad | www.encodeproject.org |
| HistMarks | ENCFF039JOT | ENCSR000DRX | ChIP-seq | H3K27me3 | John Stamatoyannopoulos, UW | www.encodeproject.org |
| HistMarks | ENCFF340JIF | ENCSR000AKC | ChIP-seq | H3K27ac | Bradley Bernstein, Broad | www.encodeproject.org |
| HistMarks | ENCFF028KBY | ENCSR000AKH | ChIP-seq | H3K9ac | Bradley Bernstein, Broad | www.encodeproject.org |
| HistMarks | ENCFF392LMQ | ENCSR000AOX | ChIP-seq | H3K9me3 | Bradley Bernstein, Broad | www.encodeproject.org |
| HistMarks | ENCFF171MDW | ENCSR000DRW | ChIP-seq | H3K36me3 | John Stamatoyannopoulos, UW | www.encodeproject.org |
| HistMarks | ENCFF601YET | ENCSR000AOV | ChIP-seq | H2AFZ | Bradley Bernstein, Broad | www.encodeproject.org |
| HistMarks | ENCFF003DXG | ENCSR057BWO | ChIP-seq | H3K4me3 | Bradley Bernstein, Broad | www.encodeproject.org |
| HistMarks | ENCFF309OEW | ENCSR000AKI | ChIP-seq | H4K20me1 | Bradley Bernstein, Broad | www.encodeproject.org |
| HistMarks | ENCFF803DJF | ENCSR000AOW | ChIP-seq | H3K79me2 | Bradley Bernstein, Broad | www.encodeproject.org |
| HistMarks | ENCFF327BUB | ENCSR702RZM | ChIP-seq | H3K27me3 | Bradley Bernstein, Broad | www.encodeproject.org |
| HistMarks | ENCFF094AZY | ENCSR117IQD | ChIP-seq | H3K9me3 | Bradley Bernstein, Broad | www.encodeproject.org |
| HistMarks | ENCFF986YKG | ENCSR887ESB | ChIP-seq | H3K4me1 | Bradley Bernstein, Broad | www.encodeproject.org |
| HistMarks | ENCFF652DFI | ENCSR580WRV | ChIP-seq | H3K36me3 | Bradley Bernstein, Broad | www.encodeproject.org |
| DHS | ENCFF598KWZ | ENCSR000EMT | DNase-seq | NA | John Stamatoyannopoulos, UW | www.encodeproject.org |
| DHS | ENCFF073ORT | ENCSR000EMT | DNase-seq | NA | John Stamatoyannopoulos, UW | www.encodeproject.org |
| DHS | ENCFF749FDY | ENCSR239XNU | DNase-seq | NA | John Stamatoyannopoulos, UW | www.encodeproject.org |
| TSAseq | SRR5516303 | PRJNA321975 | TSA-seq | SON - sample | Belmont, UIUC | www.ebi.ac.uk/ena/ |
| TSAseq | SRR5516304 | PRJNA321975 | TSA-seq | SON - control | Belmont, UIUC | www.ebi.ac.uk/ena/ |
| TSAseq | SRR7065754 | PRJNA321975 | TSA- | Lamin A/C - | Belmont, UIUC | www.ebi.ac.uk/ena/ |

| Dataset type | file accession | experiment/ project accession | assay name | target | lab | data availability |
|---|---|---|---|---|---|---|
| | | | seq | sample | | |
| TSAseq | SRR7065755 | PRJNA321975 | TSA-seq | Lamin A/C - control | Belmont, UIUC | www.ebi.ac.uk/ena/ |
| TSAseq | SRR7065760 | PRJNA321975 | TSA-seq | Lamin B - sample | Belmont, UIUC | www.ebi.ac.uk/ena/ |
| TSAseq | SRR7065761 | PRJNA321975 | TSA-seq | Lamin B - control | Belmont, UIUC | www.ebi.ac.uk/ena/ |
| LADs | GSE94971 | PRJNA374977 | DamID-seq | Lamin B1 | Schirmer, U. of Edinburgh | www.ncbi.nlm.nih.gov/geo |
| | ENCFF356LFX | ENCSR636HFF | | exclusion list regions | Anshul Kundaje, Stanford | www.encodeproject.org |

**Table S2** - Genomic annotation datasets used in this work

| Predictor | Parameter Estimates | Std. Error | t value | $-log_{10}(p.value)$ | $R^2$ | adjusted $R^2$ |
|---|---|---|---|---|---|---|
| Local host expression intensity | -0.08 | 0.00 | -28.44 | 154.97 | 0.23 | 0.23 |
| Proximity to nuclear speckle | -0.38 | 0.01 | -28.17 | 152.36 | 0.23 | 0.23 |
| GC levels | -2.14 | 0.08 | -26.99 | 141.34 | 0.21 | 0.21 |
| Gene density | -0.27 | 0.01 | -23.15 | 107.47 | 0.17 | 0.17 |
| Distance from centromere (Gb) | -4.92 | 0.28 | -17.75 | 66.02 | 0.11 | 0.11 |

**Table S3** - Univariate linear regression, potential predictors of HTLV-1 survival

| Predictor | Parameter Estimates | Std. Error | t value | $-log_{10}(p.value)$ | $R^2$ | adjusted $R^2$ |
|---|---|---|---|---|---|---|
| Proximity to nuclear speckle | -0.32 | 0.03 | -11.89 | 30.87 | 0.06 | 0.06 |
| GC levels | -1.83 | 0.16 | -11.65 | 29.73 | 0.06 | 0.06 |
| Gene density | -0.23 | 0.02 | -9.48 | 20.18 | 0.04 | 0.04 |
| Local host expression intensity | -0.06 | 0.01 | -8.21 | 15.41 | 0.03 | 0.03 |
| Distance from centromere (Gb) | -1.44 | 0.52 | -2.79 | 2.27 | 0.00 | 0.00 |

**Table S4** - Univariate linear regression, potential predictors of HIV-1 survival
